# The Rab32-LRMDA-Retriever Complex is a Key Regulator of Intestinal Immune Homeostasis

**DOI:** 10.1101/2025.07.16.665158

**Authors:** Ran Song, Chigozie Ngoka, Amika Singla, Daniel A. Kramer, Alka Diwaker, Daniel J. Boesch, Qi Liu, James J. Moresco, Bruce Beutler, Hans-Christian Reinecker, Ezra Burstein, Baoyu Chen, Emre E. Turer

**Author notes:** Equal contributions (Ran Song, Chigozie Ngoka).

## Abstract

Maintaining intestinal homeostasis relies on the intricate interplay among the mucosal epithelium, immune system, and host microbiome. A key question is how innate immune cells sense and process microbes in the gut lumen, eliciting appropriate protective responses without causing tissue injury. Clearance of invading microbes and initiation of downstream inflammatory responses are central to this process and require proper function of the endolysosomal system. Dysfunction of this system can predispose the host to chronic inflammatory disorders and acute infections. Here, through forward genetic screening of N-ethyl-N-nitrosourea (ENU)-mutagenized mice and CRISPR/Cas9 validation, we identify *Lrmda*, encoding leucine-rich melanocyte differentiation-associated protein (LRMDA), as a key regulator of intestinal homeostasis. Using hematopoietic chimera and conditional knockouts, we show that LRMDA functions primarily in CD11c^+^ cells, including mucosal dendritic cells (DCs) and macrophages, but not in non-hematopoietic cells. Proteomic, cellular, and biochemical analyses reveal that LRMDA directly and cooperatively interacts with the endolysosome-specific small GTPase Rab32 and the endosomal recycling complex Retriever. Loss of LRMDA or Retriever function increases susceptibility to dextran sodium sulfate (DSS)-induced colitis and impairs clearance of *Listeria monocytogenes*. Together, our findings establish the Rab32-LRMDA-Retriever complex as a critical regulator of endolysosomal trafficking in innate immune cells, essential for maintaining intestinal immune homeostasis.

## Introduction

Intestinal homeostasis depends on complex interactions among multiple cellular components of the intestinal mucosa, such as epithelial and immune cells, where they interact with the high burden of microorganisms in the intestinal lumen. Failure to maintain intestinal homeostasis increases susceptibility to gastrointestinal infections and chronic inflammation such as ulcerative colitis and Crohn’s disease.^1–3^ In the lamina propria, myeloid cells such as dendritic cells (DCs) and mucosal macrophages play key roles by clearing pathogens and regulating immune responses^4^. However, the molecular mechanisms through which these innate immune cells maintain intestinal homeostasis remain incompletely understood. Multiple studies have indicated that intracellular trafficking through the endolysosomal system is pivotal to myeloid cell function. This dynamic network of vesicles, which also includes phagosomes in myeloid cells, maintain constant flux with the plasma membrane, lysosomes, and lysosome-related organelles (LROs), which share core features with lysosomes but carry variable cargos depending on the cell type. Proper endolysosomal trafficking ensures that various phagocytic receptors, ion channels, transporters, effector molecules, and internalized microorganisms are trafficked to the proper cellular destination, which is essential for microbial clearance.^5–7^ In fact, many microbial effector proteins target endolysosomal trafficking as an escape mechanism to promote the intracellular survival of pathogens.^8–11^

Precise control of endolysosomal trafficking relies on the coordinated action of many proteins, among which the cargo recycling complex Retriever plays a pivotal role. Retriever consists of three subunits (VPS35L, VPS29, and VPS26C) and can dynamically associate with the CCC complex (comprising CCDC22, CCDC93, DENND10, and ten COMMD proteins COMMD1-10) to form a larger supramolecular assembly known as the Commander complex.^12–14^ Largely through its interaction with the cargo adaptor sorting nexin 17 (SNX17), Retriever mediates the recycling of cargo proteins critical for innate immunity from endosomes to the plasma membrane, including various integrins, tyrosine receptor kinases, G-protein coupled receptors (GPCRs), and lipoprotein receptors.^13^ Mutations in Retriever or CCC components lead to Ritscher-Schinzel syndrome, a disorder characterized by developmental anomalies.^15–17^ The role of Retriever in immune cells and its contributions to innate immunity remain largely unknown.

To identify key regulators of gastrointestinal homeostasis, we performed a forward genetic screen using N-ethyl-N-nitrosourea (ENU)-mutagenized mice and assessed their susceptibility to dextran sodium sulfate (DSS)-induced colitis, a well-established model of epithelial damage that triggers gastrointestinal inflammation.^18, 19^ Through this screen, we identified two independent mutations in *Lrmda* resulting in severe colitis. In humans, missense and nonsense mutations in LRMDA (also known as Oca7 or C10orf11) are associated with oculocutaneous albinism, in which LRMDA is implicated in the biogenesis of melanosomes, an specialized LRO akin to those found in innate immune cells.^20^ However, the molecular mechanisms by which LRMDA regulates LRO biogenesis or mucosal inflammation remain unexplored. We find that LRMDA directly and cooperatively interacts with the LRO-specific small GTPase Rab32 and with Retriever. These interactions are specifically disrupted by disease-associated mutations. Furthermore, we show that both LRMDA and Retriever are essential for containing *Listeria monocytogenes* within Rab32^+^ vesicles and facilitating its subsequent clearance. Together, our studies uncover the Rab32-LRMDA-Retriever complex as an essential regulator of endolysosomal trafficking in innate immune cells and a key modulator intestinal immune response.

## Results

### LRMDA deficiency leads to colitis susceptibility

To identify genes critical for intestinal immune regulation, we performed random mutagenesis using N-ethyl-N-nitrosourea (ENU) and applied a previously described inbreeding scheme to assess both heterozygous and homozygous mutant phenotypes.^18, 21–23^ Mice were screened for susceptibility to dextran sodium sulfate (DSS)-induced colitis by monitoring daily weight loss. This screen identified two independent ENU pedigrees (named *bowie* and *stardust* for the late British rock star) in which mutations—both in the coding sequence of *Lrmda*—were strongly associated with enhanced weight loss and colitis susceptibility in an additive model of inheritance (p-value 2.2 × 10^-12^) (**Fig. 1a-c**). The *bowie* mutation introduced a premature stop in exon 1 (Y13*), while the *stardust* mutation encoded a D37A missense substitution, which was predicted to be damaging by PolyPhen-2 (score = 0.999)^24^. AlphaFold3 modeling indicated that LRMDA comprises a well-folded amino-terminal (NT) leucine-rich repeat (LRR) domain followed by an unstructured carboxy-terminal (CT) tail (**Fig. 1d**).^25^ The D37 residue lies on the third beta strand in the LRR domain, where it forms multiple polar interactions with the NH of S39 and the side chains of S39, S12, and R35, suggesting a potential role in stabilizing the LRR fold (**Fig. 1d, e**).^26^ Consistent with this prediction, the D37A mutation abolished LRMDA expression in virally transduced myeloid cells (**Fig. 1f**) and impaired protein solubility in *E. coli* overexpression, even when fused to the solubility-enhancing tag maltose binding protein (MBP) (**Fig. 1g**). Together, these data indicate that loss-of-function mutations in *Lrmda* compromise protein stability and render mice susceptible to DSS-induced colitis.

**Figure 1.**
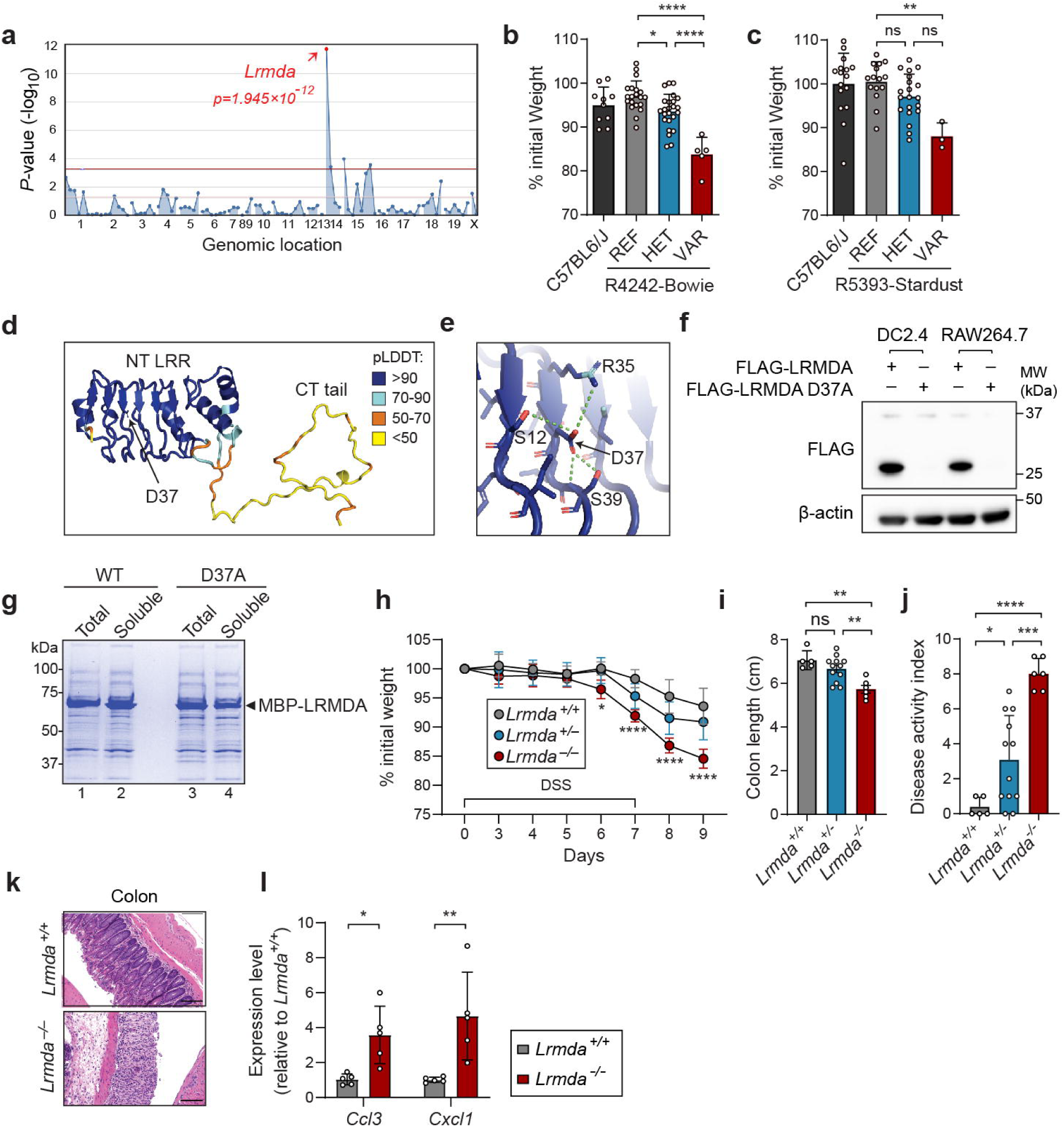
Mapping and validation of *Lrmda* in *Bowie* and *Stardust* phenotype. **(a)** Manhattan plot showing *P* values of association between *stardust* and mutations identified in *stardust* pedigree, calculated using an additive model of inheritance. The −log_10_ *P* values (*y*-axis) are plotted versus the chromosomal positions of the mutations (*x*-axis). Horizontal red and purple lines indicate thresholds of *P*=0.05 with or without Bonferroni correction, respectively. *P* values for linkage of mutation in *Lrmda* with the *stardust* DSS phenotype are indicated. **(b, c)** Percentage of body weight loss on day 10 of DSS treatment for each genotype. (b) Data are shown for C57BL6/J (n=10), Reference (REF) *Lrmda^+/+^*(n=20), Heterozygous (HET) *Lrmda^Bowie/+^* (n=24), and Variant (VAR) *Lrmda^Bowie/Bowie^*(n=5) in the R4242 pedigree (\**P*=0.0197, \*\*\*\**P*<0.0001). (c) Data are shown for C57BL6/J (n=16); REF *Lrmda^+/+^* (n=15); HET *Lrmda^Stardust/+^*(n=20); VAR *Lrmda^Stardust/Stardust^* (n=3) in the R5393 pedigree (\**P*=0.0042). **(d)** AlphaFold3-predicted structural model of LRMDA, featuring an N-terminal leucine-rich repeat (LRR) domain and an unstructured C-terminal tail. Structure is colored using predicted local difference distance test (pLDDT) scores, with high scores indicate high reliability in local structure prediction. Position of aspartate 37 (D37) is indicated. **(e)** Close-up view of structures surrounding D37. Polar interactions are indicated with dashed green lines. **(f)** Immunoblot of lysates from DC2.4 cells and RAW264.7 cells transduced with wild-type (WT) or D37 mutant LRMDA. β-actin was used as a loading control. **(g)** Coomassie-blue stained SDS PAGE gel comparing solubility of MBP-tagged LRMDA expressed from *E. coli* cells. Total and soluble protein fractions are shown. **(h)** Weight loss analysis of CRISPR/Cas9-targeted *Lrmda^+/+^*, *Lrmda^+/-^*, and *Lrmda^-/-^* mice following 1.4% DSS treatment (\**P*=0.0131, \*\*\*\**P*<0.0001). **(i, j)** Colonic length (\*\**P*=0.0012, \*\**P*=0.0048) (i) and disease activity index (\**P*=0.0448, \*\*\**P*=0.0002, \*\*\*\**P*<0.0001) (j) after 10 days of DSS treatment in *Lrmda^+/+^*, *Lrmda^+/-^*, and *Lrmda^-/-^* mice. **(k)** Representative H&E-stained colon sections from *Lrmda^+/+^* and *Lrmda^-/-^*mice following DSS treatment. Scale bars: 50 μm. **(l)** Quantitative PCR analysis of distal colon tissue from DSS-treated *Lrmda^+/+^* and *Lrmda^-/-^* mice. Expression levels are presented as fold change relative to *Lrmda^+/+^* controls (\**P*=0.0341, \*\**P*=0.0030). Data are expressed as means ± s.d. Statistical significance was determined by one-way analysis of variance (ANOVA) with Turkey’s multiple comparisons (b, c, i, j), two-way ANOVA with Turkey’s multiple comparisons (h), or two-way ANOVA with Šídák’s multiple comparisons (l). ns, not significant.

To confirm that *Lrmda* is the causative gene in these colitis-susceptible pedigrees, we used CRISPR/Cas9 gene editing to introduce a null mutation in the second coding exon (**Extended Data Fig. 1**). Consistent with the ENU-induced mutations, the resulting *Lrmda^-/-^* mice exhibited significantly greater weight loss in response to DSS treatment compared to the wild-type and *Lrmda^+/-^* controls (**Fig. 1h**). Additionally, *Lrmda^-/-^* mice displayed shortened colons, increased disease activity index (DAI), and more severe histologic features of colitis compared to wild-type or *Lrmda^+/-^* mice (**Fig. 1i-k**). The distal colons of *Lrmda^-/-^* mice also showed increased mRNA expression of the proinflammatory chemokines CCL3 and CXCL1 (**Fig. 1l**). Overall, these data confirm that loss of *Lrmda* function is the cause of increased susceptibility to colitis.

### LRMDA functions within the innate immune compartment

To determine the cellular context in which LRMDA functions, we analyzed single-cell RNA-seq datasets from human patients with ulcerative colitis and Crohn’s disease.^27, 28^ Within the immune compartment, *Lrmda* transcripts were enriched in dendritic cells, macrophages, and monocytes, with lower expression in mast cells (**Fig. 2a, Extended Data Fig. 2a**). No expression was detected in the stroma or epithelial layers of the intestine. In human Crohn’s disease and ulcerative colitis samples, an increase in *Lrmda* transcript is found in the ileum and colon, which likely reflects an influx of inflammatory cells into these tissues during these disease processes (**Extended Data Fig. 2b, c**).^29^ Assessing the peripheral and lamina propria immune compartments of *Lrmda^-/-^* animals revealed no differences in major immune subsets, arguing against a developmental defect in immune cell populations (**Extended Data Fig. 3**). These expression data support a model in which LRMDA mainly functions within the innate immune compartment to mitigate intestinal inflammation, rather than acting through the epithelial or stromal compartments.

**Figure 2.**
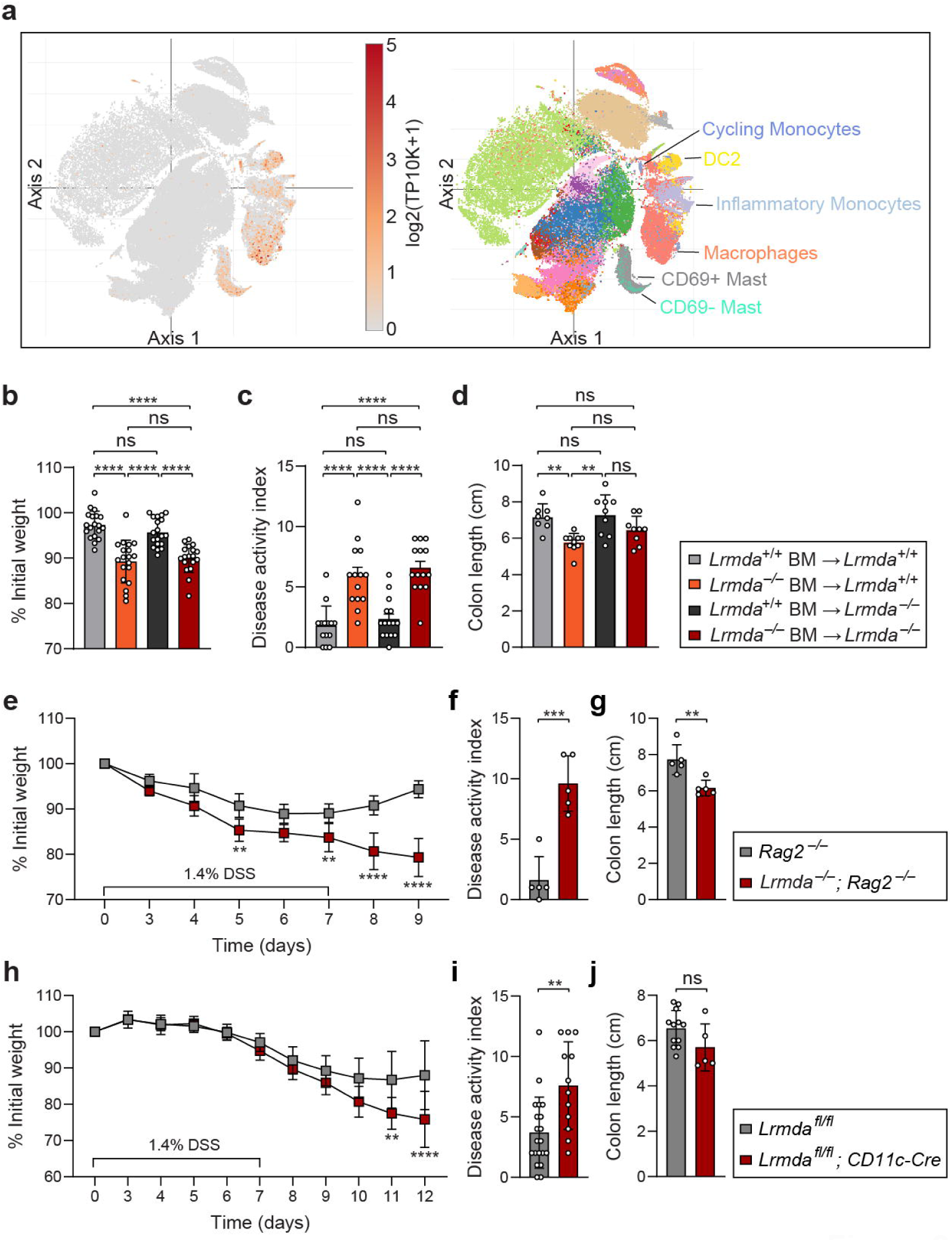
Expression and function of LRMDA in innate immune cells. **(a)** UMAP of single-cell RNA-seq data showing *Lrmda* expression in samples from ulcerative colitis and Crohn’s disease patients. **(b-d)** Weight change (\*\*\*\**P*<0.0001) (b), disease activity index (\*\*\*\**P*<0.0001) (c), and colon length (\*\**P*=0.0045, \*\**P*=0.0015) (d) of bone marrow chimeras treated with 1.4% DSS. **(e-g)** Weight loss (\*\**P*=0.0061, \*\**P*=0.0069, \*\*\*\**P*<0.0001) (e), disease activity index (\*\*\**P*=0.0003) (f), and colon length (\*\**P*=0.0057) (g) of *Lrmda*^-/-^; *Rag2*^-/-^ mice treated with 1.4% DSS. **(h-j)** Weight loss (\*\**P*=0.0011, \*\*\*\**P*<0.0001) (h), disease activity index (\*\**P*=0.0024) (i), and colon length (j) of CD11c⁺ myeloid-specific *Lrmda*-deficient mice (*Lrmda^fl/fl^; CD11c-Cre*) after DSS treatment. Data are expressed as means ± s.d. Statistical significance was determined by one-way ANOVA with Turkey’s multiple comparisons (b, c, d), two-way ANOVA with Turkey’s multiple comparisons (e, h), or unpaired Student t-test (f, g, i, j). ns, not significant.

To test this model, we first asked whether the colitis phenotype results specifically from the loss of LRMDA in hematopoietic cells. For this, we generated bone marrow chimeric mice by irradiating 8-10-week-old recipients and transplanting bone marrow from *Lrmda^-/-^*or wide-type littermate controls. After DSS administration, chimeric mice transplanted with *Lrmda^-/-^* hematopoietic cells exhibited significant weight loss and increased susceptibility to colitis, irrespective of host genotype (**Fig. 2b**). In contrast, *Lrmda^-/-^* mice transplanted with wild-type bone marrow were protected and displayed weight loss similar to wild-type control animals (**Fig. 2b**). The weight loss correlated with increased rectal bleeding and diarrhea, as well as reduced colon length (**Fig. 2c, d**). These data confirm that LRMDA functions in hematopoietic cells to protect against colitis.

To directly assess the role of LRMDA in innate immune cells, we employed two complementary approaches: 1) generating double knockout mice lacking *Lrmda* and *Rag2*, and creating a conditional allele in which *Lrmda* is only deleted in CD11c^+^ cells. In the first model, *Lrmda^-/-^; Rag2^-/-^* animals had significantly greater weight loss and DAI compared to *Rag2^-/-^* controls following DSS treatment (**Fig. 2e-g**), demonstrating that the adaptive immune system is dispensable for the LRMDA-dependent protection against colitis. In the second model, we crossed mice with a floxed allele of *Lrmda* (with LoxP sites flanking exon 3) to the Itgax-Cre (CD11c-Cre) background, which targets CD11c^+^ mucosal dendritic cells and macrophages. The resulting cell-specific knockout mice displayed increased sensitivity to colitis, showing greater weight loss and increased DAI upon DSS administration (**Fig. 2h-j**). Together, these data establish that LRMDA functions within the myeloid cell compartment to prevent colitis and maintain intestinal homeostasis, independent of antigen presentation by the adaptive immune system.

### LRMDA directly interacts with Retriever and Rab32

To gain mechanistic insights into LRMDA function, we performed unbiased proteomic analysis to define its interactome. For this, we established stable cell lines expressing FLAG-tagged LRMDA through lentiviral transduction in both dendritic (DC2.4) and macrophage (RAW264.7) cell lines. Mass spectrometry analysis revealed 14 proteins associated with LRMDA across both cell types; among them, 13 proteins were components of the Retriever or CCC complex (**Fig. 3a**). In addition, we also detected Rab32 in DC2.4 cells (**Supplemental Table 1**), consistent with recent yeast two-hybrid studies identifying an LRMDA-Rab32 interaction.^20^

**Figure 3.**
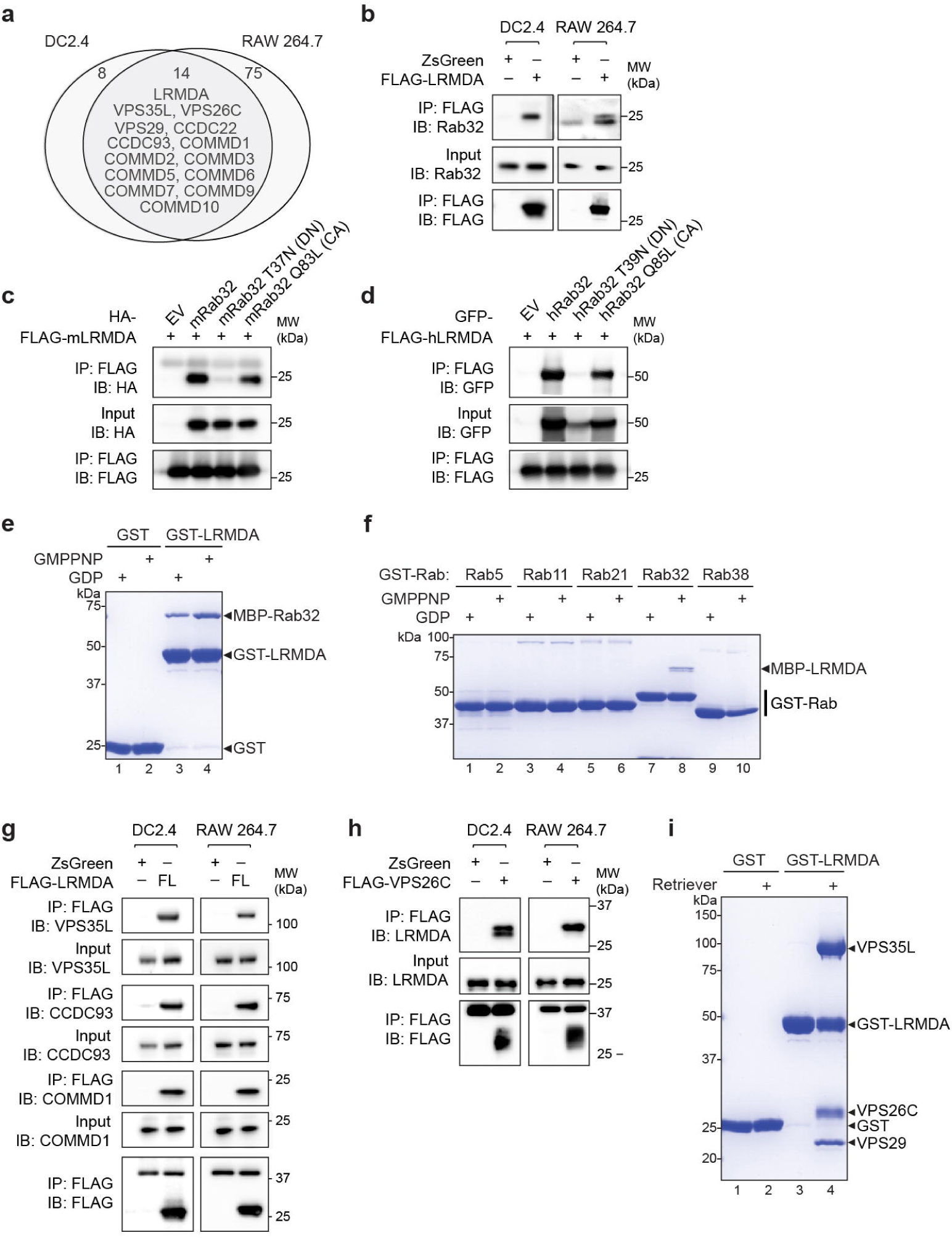
Identification of Retriever and Rab32 as LRMDA-interacting proteins. **(a)** Venn diagram of LRMDA-interacting proteins in DC2.4 and RAW264.7 cells transduced with FLAG-tagged LRMDA, identified by anti-FLAG IP followed by mass spectrometry. **(b)** Immunoblot using anti-Rab32 antibody following FLAG-IP from FLAG-LRMDA transduced DC2.4 and RAW264.7 cells. MW, molecular weight; IB, immunoblot. **(c-d)** Co-IP and immunoblot of mouse (c) and human (d) LRMDA with WT or mutant Rab32 in HEK293T cells. **(e)** Coomassie blue-stained SDS–PAGE gels showing GST-LRMDA pull-down of MBP-Rab32 loaded with GDP vs. the nonhydrolyzable GTP analogue GMPPNP. **(f)** Coomassie blue-stained SDS-PAGE gel of indicated GST-tagged Rab proteins pulling down MBP-LRMDA in different nucleotide states. **(g)** IP and immunoblot for VPS35L, CCDC93, and COMMD1 from FLAG-LRMDA transduced cells. **(h)** IP and immunoblot for LRMDA from cells transduced with FLAG-tagged VPS26C. **(i)** Coomassie blue-stained SDS–PAGE gel showing the pull-down of purified Retriever by GST-LRMDA.

First, we validated the interaction between LRMDA and Rab32 using co-immunoprecipitation (co-IP) in stable DC2.4 and RAW264.7 cell lines expressing FLAG-LRMDA. In both cell types, FLAG-LRMDA precipitated Rab32 (**Fig. 3b**). When transfected in HEK293T cells, both human and mouse Rab32 bind to LRMDA in an activity-dependent manner, with the dominant negative (DN) mutants of Rab32 showing minimal binding, while the constitutively active (CA) forms maintaining robust binding (**Fig. 3c, d**). We further confirmed that this interaction is direct and prefers the GTP-bound form of Rab32, as shown in GST pull-down assays using purified recombinant proteins (**Fig. 3e lane 3-4, 3f lane 7-8**). Notably, this interaction was specific to Rab32 and was not observed for other tested Rab GTPases, including a closely related Rab protein, Rab38 (**Fig. 3f**). ^30^

We next validated the interaction between LRMDA and Retriever following the same strategy. In both DC2.4 and RAW264.7 cell lines, FLAG-LRMDA robustly precipitated endogenous Retriever (VPS35L) and the associated CCC complex (represented by CCDC93 and COMMD1) (**Fig. 3g**). A reciprocal co-IP using FLAG-tagged VPS26C also specifically precipitated endogenous LRMDA (**Fig. 3h**). Furthermore, GST pull-down analyses using purified recombinant proteins confirmed that the interaction between LRMDA and Retriever is direct (**Fig. 3i**). Together, these data demonstrate that LRMDA directly interacts with both the LRO-specific GTPase Rab32 and the endosomal recycling complex Retriever.

### Retriever deficiency phenocopies LRMDA deficiency

Given that Retriever and Rab32 are the major interactors of LRMDA, we hypothesized that these molecules function in the same genetic pathway as LRMDA to prevent colitis and maintain intestinal homeostasis. Consistent with this idea, previous studies have established that loss of Rab32 in mucosal dendritic cells and macrophages increases susceptibility to DSS-induced colitis.^31^ However, whether Retriever contributes to this pathway has remained unknown. To address this, we generated conditional *Vps35l*-deficient mice by flanking exon 11 of the gene with LoxP sequences and breeding them with *Ubc^Cre-ERT^*^2^ animals. Offspring were treated with tamoxifen to induce *Vps35l* gene deletion in adulthood (**Fig. 4a**). Upon DSS treatment, *Vps35^fl/fl^*, *Ubc^Cre-ERT^*^2^ mice displayed significantly increased susceptibility to colitis, with peak weight loss reaching 20% compared to 5% in littermate controls (**Fig. 4b**). Moreover, these mice also displayed increased colonic mRNA levels of the inflammatory cytokines *Cxcl10* and *Tnf* (**Fig. 4c**). These findings demonstrate that Retriever, like LRMDA, plays a critical role in protecting against intestinal inflammation and maintaining mucosal homeostasis.

**Figure 4.**
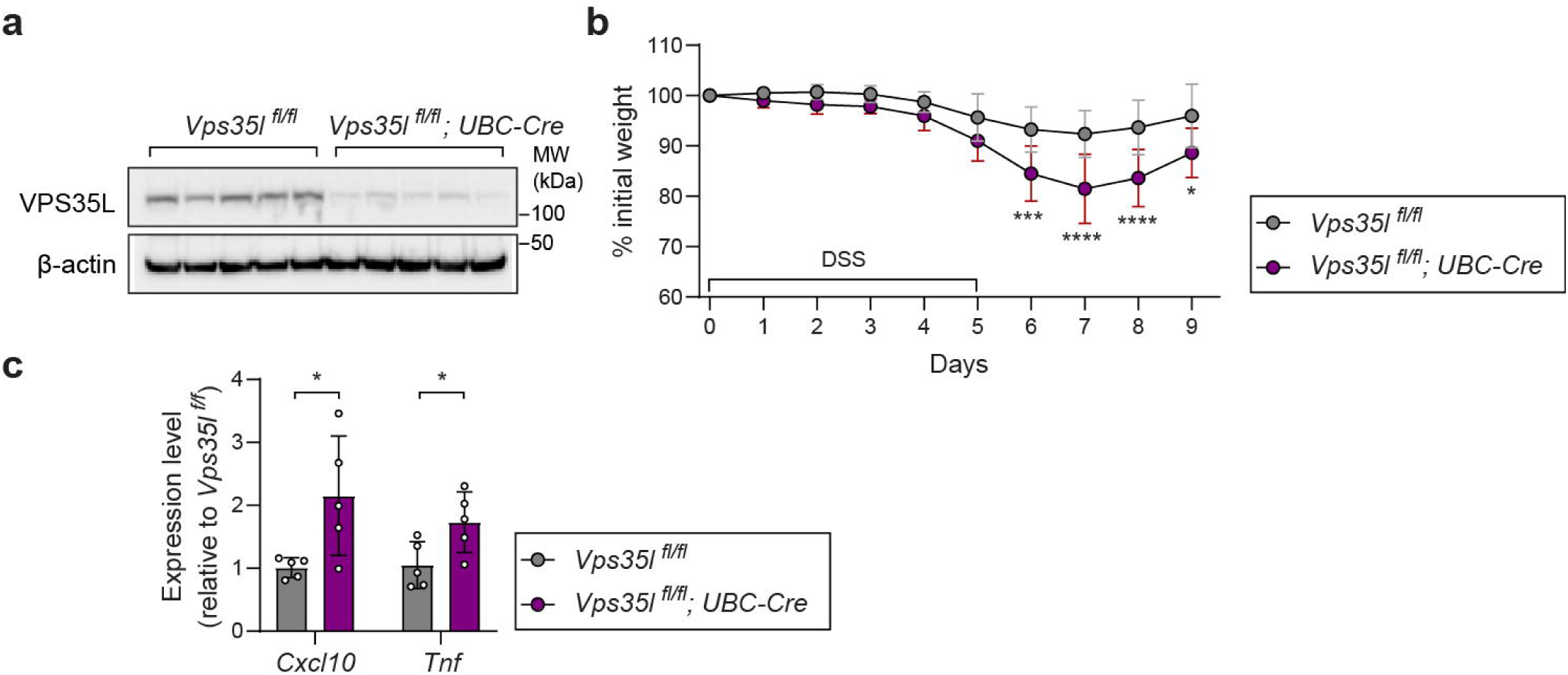
Retriever deficiency similarly increases sensitivity to DSS-induced colitis. **(a)** Immunoblot of colon tissue from tamoxifen-treated *Vps35l*^fl/fl^ and *Vps35l*^fl/fl^*; UBC-Cre* mice. **(b)** Body weight loss in tamoxifen-treated *Vps35l*^fl/fl^ and *Vps35l*^fl/fl^*; UBC-Cre* mice after 2.5% DSS treatment (\**P*=0.0161, \*\*\**P*=0.0001, \*\*\*\**P*<0.0001) **(c)** qPCR analysis of distal colon tissue from DSS-treated *Vps35l*^fl/fl^ and *Vps35l*^fl/fl^*; UBC-Cre* mice following tamoxifen treatment. Expression levels are presented as fold change relative to *Vps35l*^fl/fl^ controls (\**P*=0.0290, \**P*=0.0366). Data are expressed as means ± s.d. Statistical significance was determined by two-way ANOVA with Turkey’s multiple comparisons (b) or unpaired Student t-test (c).

### LRMDA binds to Retriever through its C-terminal tail

To elucidate how LRMDA interacts with Retriever, we used AlphaFold3 to model the LRMDA-Retriever complex (**Fig. 5a-c**). The high-confidence model predicts that the CT tail of LRMDA, but not the NT LRR domain, mediates Retriever binding through three distinct contact points: 1) residues 165-171 at the start of the CT tail engage the N-terminal beta strands of VPS26C’s arrestin fold; 2) residues 198-206 contact the C-terminal beta strands of VPS26C; and the extreme C-terminal tip (residues 215-226) inserts into a pocket formed jointly by VPS26C and VPS35L. Notably, the binding mode and sequence of the C-terminal tip aligns with that of the recently resolved cryo-EM structure of SNX17 bound to Retriever (**Fig. 5b**). This shared binding pocket has been shown to bind C-terminal peptides containing a conserved Retriever Interacting C-terminal Tail (RICT) motif, [ILV]-x-[DEQN]-x-x-L, where “x” is any residue.^32^

**Figure 5.**
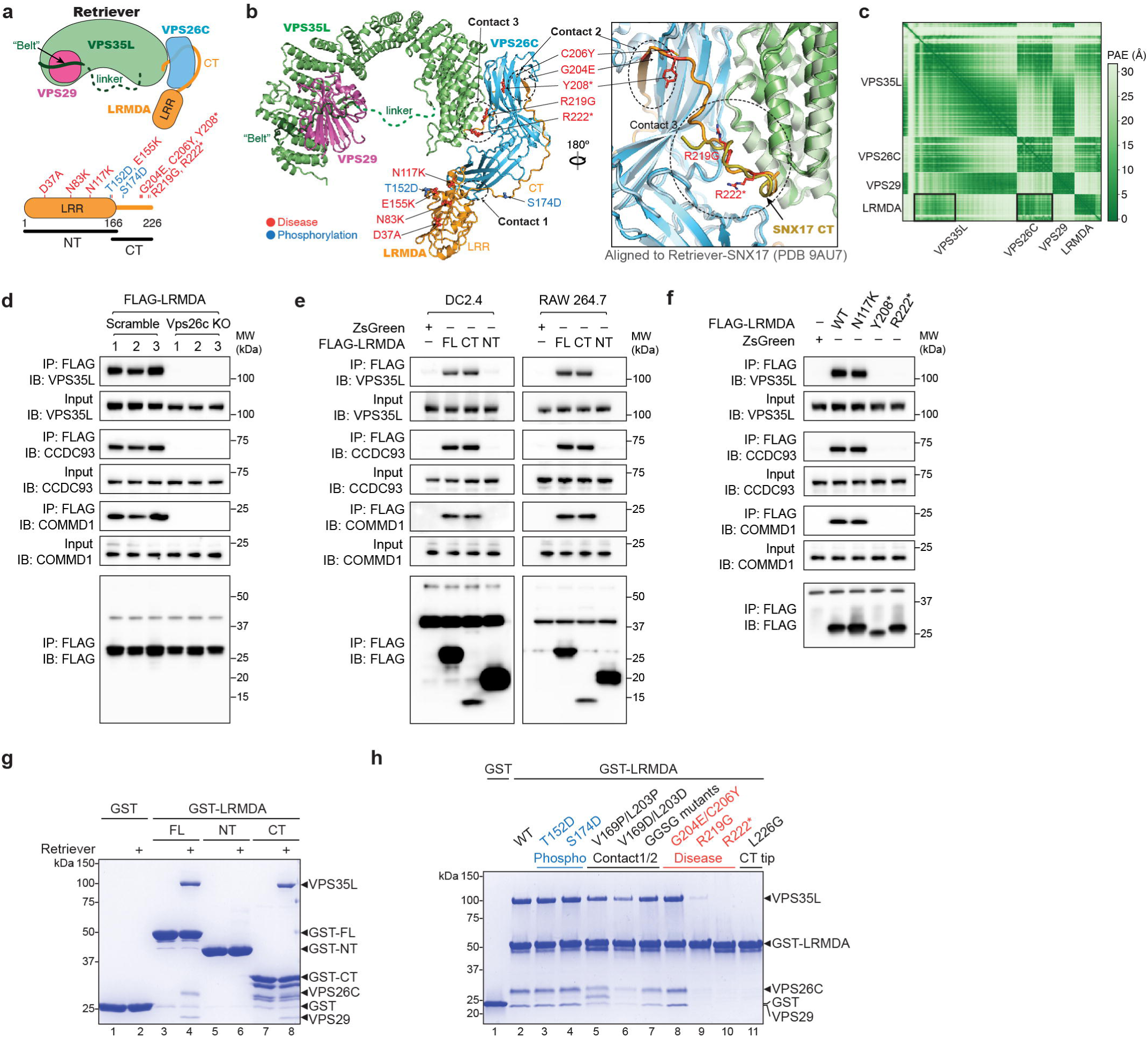
LRMDA interacts with Retriever via its C-terminal tail. **(a)** Cartoon depiction of the LRMDA-Retriever complex and domain organization of LRMDA. Disease-associated mutations (red) and phosphorylation sites (blue) are indicated. **(b)** AlphaFold3 model of the LRMDA-Retriever complex. Three predicted contacts sites are indicated by dotted circles. Disease-associated mutations (red) and phosphomimic mutations (blue) are indicated. In the close-up view, published cryo-EM structure of Retriever (grey)-SNX17(gold) complex (PDB 9AU7) is overlayed. **(c)** Predicted aligned error (PAE) score matrix of the AlphaFold3 model shown in (b), with deeper color indicating higher reliability of relative positioning of pairs of structures. Black boxes indicate contact sites of LRMDA with VPS26C and VPS35L. **(d)** IP of FLAG-LRMDA and immunoblot of endogenous Retriever complex components from Scramble or Vps26c knockout (KO) DC2.4 cells. **(e)** IP from cells transduced with ZsGreen, full-length (FL), N-terminal (NT), or C-terminal (CT) FLAG-tagged murine LRMDA and immunoblotted for endogenous Retriever and CCC proteins. **(f)** IP of FLAG-tagged human LRMDA mutants (N117K, Y208*, R222*) transduced into THP-1 cells and immunoblot of endogenous Retriever and CCC proteins (VPS35L, CCDC93, COMMD1). **(g-h)** Coomassie blue-stained SDS-PAGE gels showing pull-down of recombinant Retriever by GST-tagged LRMDA truncations (h) or full length LRMDA mutants, including disease-associated (red) and phosphomimic ones (blue)

We validated this model through several complementary approaches. First, we generated VPS26C-deficient DC2.4 cells using CRISPR/Cas9 and found that the interaction between LRMDA and Retriever/CCC complexes requires VPS26C, consistent with the participation of VPS26C in creating the binding pocket for the RICT peptide, also previously shown for other RICT containing proteins such as SNX17 and SNX31 (**Fig. 5d**).^32^ Second, we found that only full-length LRMDA or its CT tail (despite its lower stability), but not its NT LRR domain, could mediate this interaction (**Fig. 5e**). We next examined disease-associated LRMDA mutations implicated in oculocutaneous albinism in humans, including one in the LRR domain (N117K) and two truncation mutations in the CT tail (Y208* and R222*) (**Fig. 5a,b,f**). After transducing human monocyte THP-1 cells with full-length LRMDA and disease-associated variants, we observed that both Y208* and R222* abolished the interaction with Retriever (VPS35L) and CCC (COMMD1 and CCDC93), while N117K preserved the interaction (**Fig. 5f**), supporting the critical role of the CT RICT motif in engaging with Retriever. GST pull-down assays with purified recombinant proteins further confirmed that full-length LRMDA or its CT tail bound to Retriever directly, but the NT LRR domain did not (**Fig. 5g**). Disease-associated mutations within the RICT motif (R219G and R222*) or mutating the last residue of the RICT motif (L226G) severely impacted Retriever binding (**Fig. 5h, lane 9-11**). In contrast, mutations in the CT tail but outside the RICT motif, including disease-associated mutations G204E/C206Y and those predicted to mediate the interactions at contact point 1 and 2 had little effect, except for V169D/L203D, which modestly reduced binding (**Fig. 5h, lane 5-8**). Finally, we tested whether phosphorylation sites in LRMDA identified in high-throughput phosphoproteomic studies^33^ affect its association with Retriever. Phosphomimic substitutions at T152 and S174 in the CT tail had no impact on Retriever binding (**Fig. 5h, lane 3-4**), suggesting that these modifications do not directly regulate this interaction. Together, these findings validate the AlphaFold3 model and establish that LRMDA engages Retriever primarily via its C-terminal RICT motif, following a mechanism analogous to the SNX17-Retriever interaction.^32^

### LRMDA Binds to Rab32 through its N-terminal LRR Domain

We further used AlphaFold3 to explore how Rab32 could interact with LRMDA. The high-confidence model suggests that while the CT tail of LRMDA engages Retriever, its NT LRR domain binds directly to Rab32, assembling a Rab32-LRMDA-Retriever supracomplex in which LRMDA acts as a bridge (**Fig. 6a-c**). Specifically, the Switch I and II motifs of Rab32 contact the concave surface formed by the beta strands of the LRR domain (**Fig. 6b, close-up view**), while the CT tail of Rab32, which is usually prenylated in cells, points away from the complex and can be readily oriented towards membrane. Consistent with this model, pull-down assays using recombinant proteins demonstrated that the NT LRR domain, but not the CT tail, directly binds to Rab32 (**Fig. 6d, e**). Notably, the isolated LRR domain showed stronger binding than the full-length LRMDA, suggesting this region may be autoinhibited within the intact protein.

**Figure 6.**
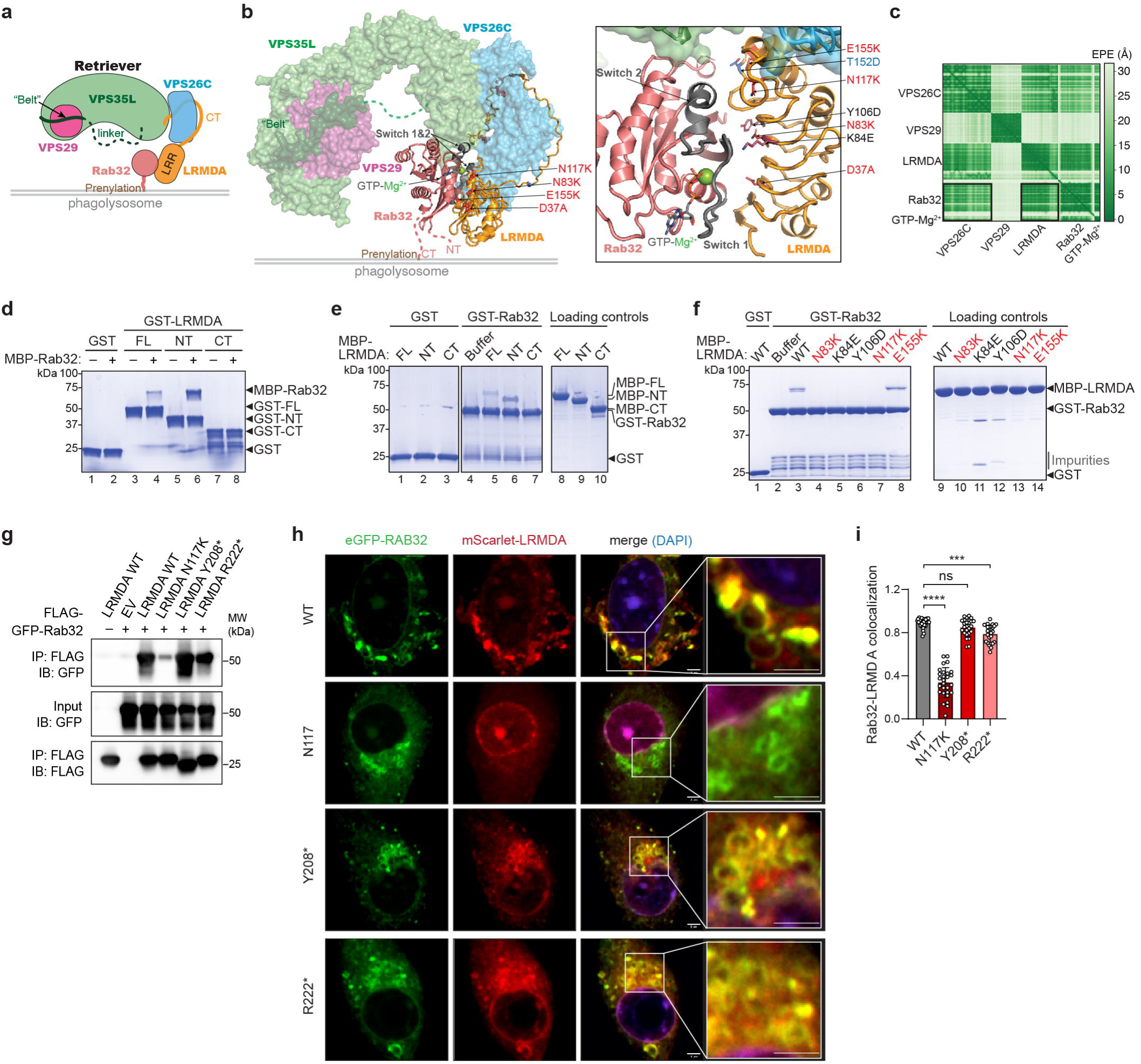
LRMDA interacts with Rab32 via its N-terminal LRR domain. **(a)** Cartoon schematic of the Rab32-LRMDA-Retriever complex, with the prenylated CT tail of Rab32 contacting phagolysosomal membranes. **(b)** AlphaFold3 model of Rab32-LRMDA-Retriever complex. Retriever is shown as semi-transparent surface models to highlight the interaction between Rab32 and LRMDA LRR domain. Disease-associated mutations (red), phosphomimic mutations (blue), and mutations designed to disrupt the interaction (black) are indicated using sticks. **(c)** PAE score matrix of the model shown in (b). Black boxes indicate direct contact of Rab32 with LRMDA and stable positioning relative to VPS26C. **(d)** Coomassie blue-stained SDS-PAGE gel showing GST-LRMDA (full length and truncations) pulling down MBP-Rab32. **(e-f)** Coomassie blue-stained SDS-PAGE gels showing GST-Rab32 pull-down of MBP-LRMDA (full length and truncations) (e) and full-length MBP-LRMDA carrying indicated mutations (f), including disease-associated ones (red). **(g)** Co-IP and immunoblot of Rab32 with wildtype or disease-associated mutant LRMDA from HEK293T cells. In (e-g), Rab32 is pre-loaded with GMPPNP. **(h-i)** Representative confocal images (h) and colocalization quantification (i) of *Lrmda* KO DC2.4 cells transduced with indicated mScarlet-tagged LRMDA and eGFP-Rab32. Nuclei were stained with DAPI.

To further validate this interaction mechanism, we tested a series of mutations within the LRR domain, including those associated with human albinism (N83K, N117K, and E155K) and two residues predicted to contact the Switch I motif of Rab32 (K84E and Y106D) (**Fig. 6b**). Consistent with the structural model, all mutations abolished Rab32 binding except E155K, which lies at the junction between the NT LRR and the CT tail and does not directly contact Rab32 (**Fig. 6f**).

We next performed co-immunoprecipitation assays using HEK293T cells transiently transfected with GFP-tagged Rab32 and selected albinism-associated LRMDA variants. As with the recombinant proteins, N117K disrupted Rab32 binding, while mutations in the C-terminal tail (Y208* and R222*) did not (**Fig. 6g**). Finally, in *Lrmda* KO DC2.4 cells transduced with mScarlet-tagged LRMDA, we observed colocalization of Rab32 and LRMDA in vesicles characteristic of endolysosomal compartments. In contrast, the N117K mutation strongly disrupted LRMDA localization to Rab32^+^ vesicles, whereas the CT mutations had negligible effects (**Fig. 6h, i**). Note that N117K did not affect Rab32 localization itself, suggesting that Rab32 recruited independently to the membrane and is responsible for subsequently recruiting LRMDA. This is consistent with the Alphafold model in which the unstructured CT tail of Rab32, which is prenylated in cells, is compatible with membrane association (**Fig. 6a, b**).

### LRMDA Cooperatively Bridges Retriever to Rab32

The AlphaFold model suggests a compelling mechanism by which LRMDA recruits Retriever to Rab32^+^ compartments by simultaneously binding both proteins to assemble a Rab32-LRMDA-Retriever complex. To test this model, we first performed pull-down assays using recombinant proteins. We observed that GST-Rab32 interacted with Retriever only in the presence of LRMDA (**Fig. 7a, lane 7 vs. 9**), confirming that LRMDA acts as a bridging factor. Intriguingly, the presence of Retriever greatly enhanced the interaction between GST-Rab32 and LRMDA (**Fig. 7a, lane 8 vs. 9**). This cooperative effect can be observed even when Rab32 was loaded with GDP, which substantially reduces its interaction with LRMDA (**Fig. 7a, lane 4 vs. 5**). Together with our observation that the isolated NT LRR domain of LRMDA binds to Rab32 more strongly than full-length LRMDA (**Fig. 6d, e**), these results strongly suggest that the LRR domain is autoinhibited in a closed conformation. Binding of Retriever to the CT tail (or truncation of the CT tail) appears to relieve this inhibition, promoting an open conformation that facilitates stable Rab32-LRMDA binding and cooperative assembly of the Rab32-LRMDA-Retriever complex (**Fig. 7b**).

**Figure 7.**
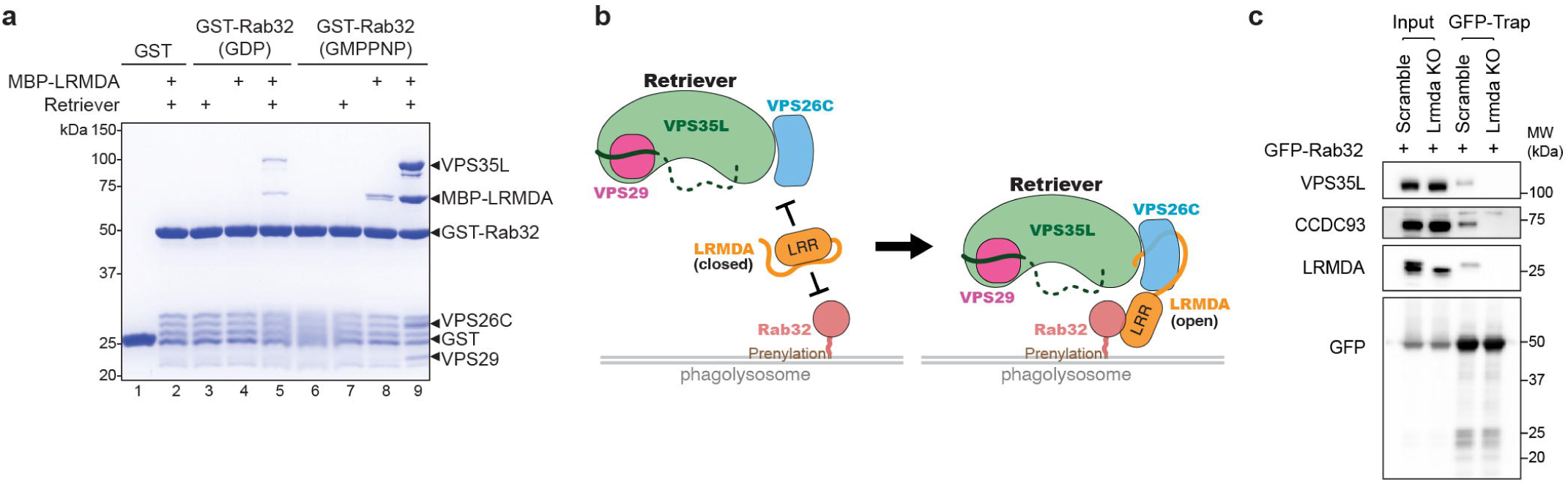
LRMDA serves as a molecular bridge linking Retriever to Rab32. **(a)** Coomassie blue-stained SDS-PAGE gel showing GST-Rab32 pull-down of recombinant Retriever complex in the absence or presence of MBP-LRMDA. **(b)** Cartoon depiction of LRMDA autoinhibition in the basal state and cooperative binding to Retriever and Rab32 in the open state, thus recruiting Retriever to Rab32^+^ membranes. **(c)** GFP-Rab32 vesicle trapping and immunoblot of Retriever components in Scramble and *Lrmda* KO DC2.4 cells transduced with GFP-Rab32.

To examine this model under more physiological conditions, we purified Rab32^+^ vesicles from cells transduced with GFP-Rab32 using GFP-Trap affinity beads under detergent-free conditions. Immunoblotting of the GFP-trapped vesicles revealed the presence of endogenous LRMDA, as well as Retriever (VPS35L) and CCC (CCDC93) (**Fig. 7c**). In contrast, Rab32^+^ vesicles purified from LRMDA-deficient DC2.4 cells lacked not only LRMDA but also Retriever and CCC (**Fig. 7c**). These findings strongly support the model that LRMDA is necessary for recruiting Retriever to Rab32^+^ vesicles.

### LRMDA and Retriever are Necessary for Bacterial Clearance

Rab32 is a pivotal GTPase associated with lysosome-related organelles (LROs), which include melanosomes essential for skin pigmentation and various specialized vesicles in immune cells critical for host defense^6, 7, 34^. In myeloid cells, Rab32 facilitates the recruitment of effector molecules to LROs required for pathogen clearance and regulation of phagolysosomal contents.^35, 36^ Rab32^+^ vesicles have been shown to be essential for eliminating intracellular bacteria such as *Mycobacterium leprae*, *Salmonella typhimurium*, and *Listeria monocytogenes*.^37–40^ However, whether LRMDA or Retriever contributes to this process has remained unknown.

We hypothesized that LRMDA and Retriever act as key effectors of Rab32 in the clearance of intracellular pathogens. To test this, we first examined the subcellular localization of *Listeria* in myeloid cells during infection. We observed that LRMDA^+^ vesicles colocalized with bacteria during *Listeria* invasion (**Fig. 8a**). In contrast, loss of LRMDA or Retriever (VPS26C) significantly reduced the recruitment of Rab32^+^ vesicles to *Listeria* (**Fig. 8b, c**). In the absence of LRMDA or Retriever, Rab32^+^ vesicles also appeared smaller and more diffusely distributed in cells (**Fig. 8b, c**). These findings suggest that LRMDA and Retriever are critical effectors that allow Rab32^+^ LROs to target bacteria effectively.

**Figure 8.**
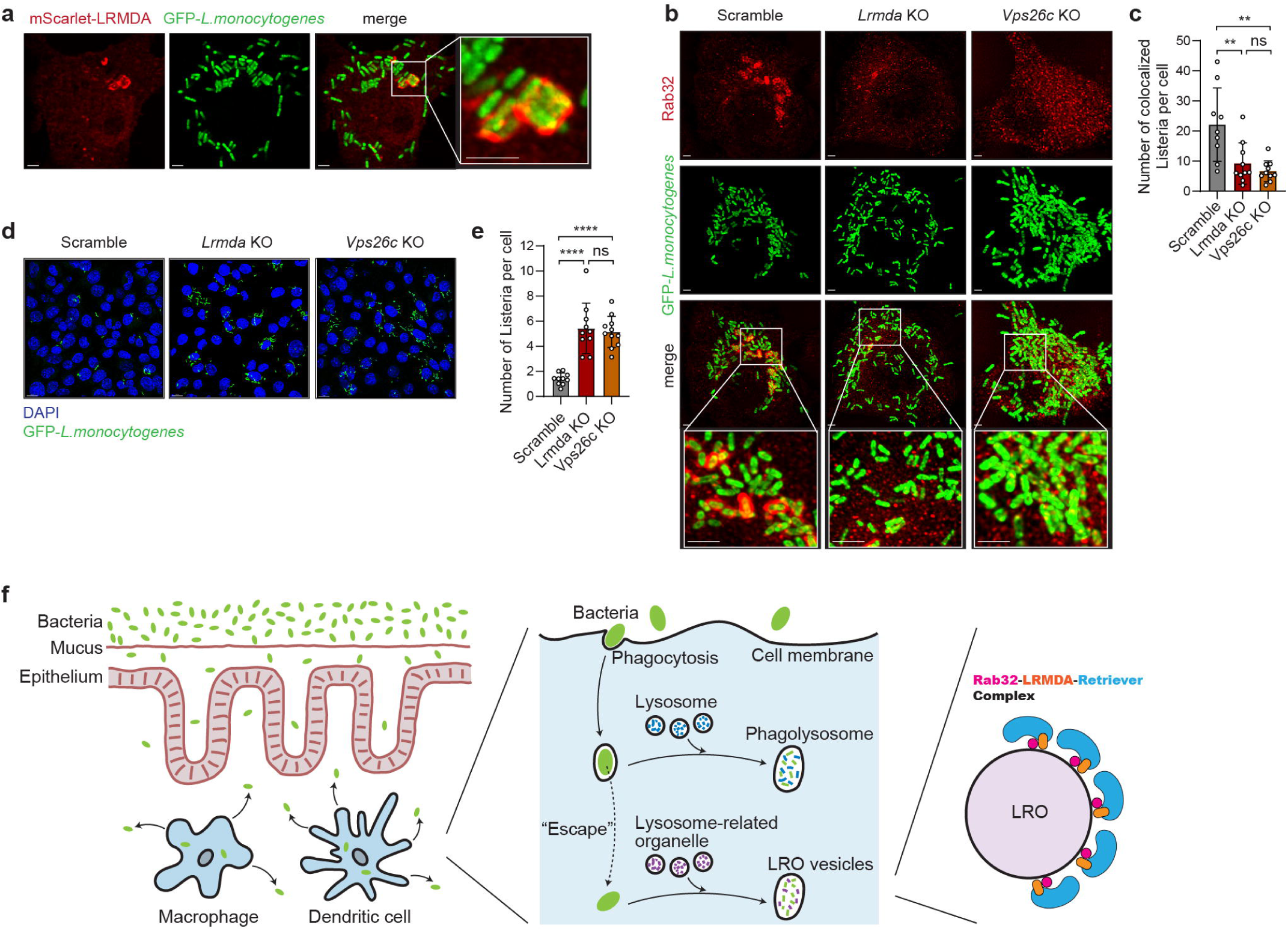
LRMDA and Retriever are required for bacterial handling and clearance. **(a)** Representative confocal images showing colocalization of LRMDA with *Listeria monocytogenes*. Scale bars: 2 μm. **(b-c)** Representative confocal images (b) and quantification of colocalization with Rab32^+^ vesicles (c) of Scramble, *Lrmda* KO or *Vps26c* KO DC2.4 cells infected with GFP-*Listeria monocytogenes* for 1.5 hours. Cells were fixed and stained with an anti-Rab32 antibody. Scale bars: 2 μm. (\*\**P*=0.0053, \*\*\**P*=0.0009). **(d-e)** Representative confocal images of DC2.4 cells (Scramble, *Lrmda* KO or *Vps26c* KO) (d) and quantification of *Listeria* (e) after GFP-*Listeria monocytogenes* infection for 1.5 hours. Nuclei were counterstained with DAPI. Scale bars: 2 μm. Data are presented as mean ± s.d. (\*\*\*\**P*<0.0001). At least 30 cells per field were analyzed per experiment. Statistical significance was determined by one-way analysis of variance (ANOVA) with Turkey’s multiple comparisons (c, e). ns, not significant. **(f)** Schematic summarizing the role of Rab32-LRMDA-Retriever complex in LRO-mediated bacterial clearance in innate immune cells and host defense against colitis.

We next assessed the impact of LRMDA and Retriever deficiency on bacterial clearance. After 90 minutes of infection, the number of uncleared *Listeria* per cell increased by more than 3-fold in cells lacking LRMDA or VPS26C (**Fig. 8d, e**). These data provide functional evidence that impaired recruitment of RAB32^+^ vesicles to pathogens compromises intracellular bacterial clearance, underscoring the essential role of the Rab32-LRMDA-Retriever axis in innate immune defense.

## Discussion

By combining forward genetics, animal colitis models, structural and mutational analyses, and pathogen clearance assays, we have identified and elucidated a critical axis in innate immune cells mediated by the Rab32-LRMDA-Retriever complex. Our findings demonstrate that, at the functional level, this complex is essential for myeloid cell function, efficient handling and clearance of invading bacteria, and host protection from colitis. At the molecular level, LRMDA functions as a bridge that cooperatively connects Retriever to Rab32^+^ vesicles.

The formation of this complex involves two distinct but cooperative interactions: the CT tail of LRMDA binding to Retriever and the NT LRR domain binding to Rab32. The CT-Retriever interaction shares the same mechanism as the Retriever-specific cargo adaptor protein SNX17 (and its ortholog SNX31), relying on the conserved RICT peptide motif at the CT tip.^12, 14^ Since the SNX17-Retriever complex is essential for recycling over a hundred membrane cargos from early endosomes back to the plasma membrane, this suggests an intriguing model in which LRMDA competes with SNX17 and redirect Retriever from early endosome to Rab32^+^ lysosome-related organelles (LROs) to regulate pathogen clearance and immune response. Note that besides SNX17, SNX31, and LRMDA, the CT RICT motif is found in other proteins, including bacterial effectors,^32^ highlighting LRMDA as an example of how Retriever can be recruited to diverse cellular contexts.

Interestingly, the LRR-Rab32 interaction is enhanced by Retriever binding to the CT or by deleting CT, suggesting that the LRR domain is autoinhibited by the CT. This provides a regulatory model in which Retriever binding to LRMDA relieves this inhibition and promotes Rab32 binding and assembly of the Rab32–LRMDA–Retriever complex. A similar autoinhibition mechanism has been observed for SNX17, where its CT tail binds to its FERM domain, blocking cargo interactions until Retriever binding frees the FERM domain to engage cargos.^32^ However, unlike SNX17, we did not observe direct binding between LRMDA’s NT LRR and CT tail. Therefore, the molecular basis of LRMDA autoinhibition remains to be determined. Several questions emerge along the same line: how is the LRMDA-Retriever interaction initiated or regulated? How do SNX17 and LRMDA coordinate hand-off of Retriever in space and time? Without SNX17, what is the precise role of Retriever in late endolysosomal vesicles? Does it mediate the sorting of distinct cargos required for their maturation and function?

Our work also provides mechanistic insights into the pathogenicity of disease-associated mutations in LRMDA. Mutations in the CT (e.g., Y208*, R219G, and R222*) disrupt the RICT motif and abolish Retriever binding, while mutations in the LRR domain (e.g., N83K and N117K) impair Rab32 binding. This suggest that the full assembly of the Rab32-LRMDA-Retriever complex is essential for their function. Conversely, other disease-associated mutations (e.g., G204E, C206Y, and E155K) and the phosphomimic substitutions (e.g., T152D and S174D) did not disrupt these interactions, suggesting the existence of additional, yet unknown regulatory mechanisms. Together, our analysis provides a roadmap for understanding the significance of LRMDA variants in innate immunity and intestinal immune homeostasis.

Previous studies established Rab32 and its guanine exchange factor (GEF) BLOC-3 (composed of Hermansky-Pudlak syndrome proteins 1 and 4) as essential regulators of LROs in various cell types, including melanocytes, platelets, macrophages, and dendritic cells.^35, 37, 40–42^ Deficiencies in these factors result in shared defects in innate immunity in humans and animal models.^36, 43, 44^ Here, we identify LRMDA and Retriever as new critical effectors of Rab32, phenocopying the effects of *Rab32* deficiency in colitis susceptibility and impaired *Listeria* handling.^37, 40, 45, 46^ Notably, LRMDA was not previously linked to immunity, and no immunologic phenotype were reported in OCA7 patients, likely due to the limited number of patients with deleterious mutations and a historical focus on ophthalmologic features.^47^ However, GWAS studies have indicated a role of LRMDA in viral responses (e.g., COVID-19), although causality has not been established.^48^ Given the link between Hermansky-Pudlak syndrome and spontaneous colitis in humans, it is plausible that careful examination of patients with *LRMDA* mutations will reveal immune and hematologic defects. Similar considerations apply to *VPS35L* deficiency, which causes 3C/Ritscher-Schinzel syndrome, a condition with severe cranio-cerebral-cardiac malformations but not yet recognized immune phenotypes, likely due to the paucity of reported cases.

The functional connection between Rab32, LRMDA and Retriever also predicts that other phenotypes seen in Rab32 deficiency will manifest with LRMDA or Retriever deficiency. For example, deficiencies in Rab32 or BLOC-3 cause skin hypopigmentation due to melanocyte dysfunction, consistent with *LRMDA* mutations causing oculocutaneous albinism.^47, 49^ Recent reports further implicate Retriever and the CCC complexes in skin pigmentation.^50^ In hematopoietic cells, other lineages also develop highly specialized and critical LROs. For example, Rab32 and BLOC-3 deficiencies are associated with platelet dysfunction, which has also been reported in animal models of CCC complex deficiency.^51^

In summary, using forward genetics, we have identified an essential and non-redundant role for *Lrmda* in maintaining intestinal homeostasis in vivo. We show that LRMDA directly binds to both the LRO regulator Rab32 and the endosome recycling complex Retriever, both of which are also necessary for pathogen clearance and intestinal immune quiescence. Our study offers direct mechanistic insights into how immune cells regulate the lysosome related organelles.

## Methods

### Mice

All animal experiments were approved by the Institutional Animal Care and Use Committee (IACUC) at the University of Texas Southwestern Medical Center and conducted in accordance with institutional guidelines. All mice were bred and housed at the University of Texas Southwestern Medical Center animal facility in accordance with institutionally approved protocols. Mice were housed at 22 °C with a 12-h light/12-h dark cycle. Animals were fed ad libitum with standard chow diet (2016 Teklad Global 16% Protein Rodent Diet) and autoclaved water.

Eight- to twelve-week-old mice were used for all experiments. All mouse experiments were performed using both male and female mice without any bias. For DSS-induced colitis, mice received 1.4% (wt/vol) DSS in drinking water for 7 days, followed by 3 days on regular water. Body weight was recorded daily and reported as a percentage of initial body weight. Disease activity index score is a composite score of weight loss, stool bleeding and stool consistency determined as previously described. Briefly: weight loss: 0, No loss. 1, 1–10% loss of body weight. 2, 10–15% loss of body weight. 3, 15–20% loss of body weight. 4, > 20% loss of body weight; stool consistency: 0 (normal), 2 (loose stool), and 4 (diarrhea); and bleeding: 0 (no blood), 1 (hemoccult positive), 2 (hemoccult positive and visual pellet bleeding), and 4 (gross bleeding and/or blood around anus).

### Generation of the Lrmda^-/-^ and conditional mouse strain using the CRISPR/Cas9 system

The generation of *Lrmda^-/-^* and conditional mouse strains by CRISPR/Cas9 used an established protocol as previously described^22^. Briefly, female C57BL/6J mice were superovulated with 6.5 U pregnant mare serum gonadotropin (Millipore), followed 48 h later by 6.5 U human chorionic gonadotropin (Sigma-Aldrich), and mated overnight with C57BL/6J males. The following day, fertilized eggs were collected from the oviducts and *in vitro*–transcribed Cas9 mRNA (50 ng/μl) and *Lrmda* small base-pairing guide RNA injected into the cytoplasm or pronucleus of the embryos. The conditional knockout allele was generated using a DNA template encoding exon 3 of *Lrmda* with flanking LoxP sites. The injected embryos were cultured in M16 medium (Sigma-Aldrich) at 37°C in 5% CO_2_. For the production of mutant mice, two-cell stage embryos were transferred into the ampulla of the oviduct (10–20 embryos per oviduct) of pseudo-pregnant Hsd:ICR (CD-1) female mice (Harlan Laboratories).

### Generation of bone marrow chimeric mice

Recipient mice were lethally irradiated with two 7-Gy exposures to X-irradiation administered 5 h apart.^22, 52^ Femurs derived from donor *Lrmda^+/+^*or *Lrmda^-/-^* mice were flushed with phosphate-buffered saline (PBS) using a 25-gauge needle. The cells were centrifuged at 700 × *g* for 5 min, and cells were resuspended in 1 mL of PBS. Bone marrow cells from *Lrmda^+/+^*or *Lrmda^-/-^* mice were transferred into the indicated recipient mice through intravenous injection. Mice received antibiotics in drinking water for 4 weeks post-transfer. DSS colitis susceptibility was assessed 8 weeks post-reconstitution.

### Cell culture, transfection, and lentiviral infection

The Lenti-X 293T cells (Takara #632180) were cultured at 37 °C in DMEM (Sigma) supplemented with10% FBS (Gibco) and 1% penicillin-streptomycin (Life Technologies) in 5% CO2. DC2.4 (Sigma-Aldrich), RAW264.7 (ATCC) and THP-1 cells (ATCC) were maintained in RPMI-1640 (Gibco) under the same supplementation. Plasmids were transfected using Lipofectamine 2000 (Life Technologies), and cells were harvested 36–48 h post-transfection. Lentivirus was produced in 293T cells using a second-generation packaging system and subsequently used to infect DC2.4, RAW264.7 and THP-1 cells.

### Plasmid

Full-length human LRMDA and Rab32 were ordered as gene strings from Thermo Fisher with NdeI/BamHI adaptors to be ligated into the pGexTev or pMalC2Tev vector for bacterial expression. LRMDA mutagenesis; V169D/L203D, G204E/C206Y, R219G, R222* (deletion of D223, D224, Q225, and L226), L226G, D37A, N83K, K84E, Y106D, N117K, D125K, D128K, A148W, E155K, F165D, LRMDA NT (1-167), LRMDA NTLong (1-177), and LRMDA CT (150-226 were created using PCR. Rab32 mutagenesis; T39N and Q85L were made using PCR. Retriever VPS35L, VPS26C, VPS29-His6 for insect cell expression were used as previously described. ^32^ A table of all plasmids used in this manuscript is available in **Supplemental Table 2**.

### Immunoprecipitation and immunoblotting

Cells were lysed with NP40 lysis buffer with glycerol (50 mM Tris–HCl pH 8.0, 150 mM NaCl, 1% (v/v) Nonidet P-40, 5% (v/v) glycerol, and protease inhibitors). Immunoprecipitation was performed using anti-FLAG M2 magnetic beads (Sigma) for 1 h at 4 °C, and the proteins were eluted with 150 μg/mL 3× Flag at 4 °C for 1.5 h. Protein concentrations were measured using a BCA assay (Pierce). Samples were loaded onto NuPAGE 4%-12% Bis-Tris protein gels (Thermo Fisher Scientific), transferred to nitrocellulose membranes (Bio-Rad), blotted with the primary antibody at 4 °C overnight and the secondary antibody for 1 h at room temperature, and then visualized by chemiluminescent substrate (Thermo Fisher Scientific).

### Quantitative reverse transcription PCR

Total RNA from colon tissue was isolated using a TRIzol reagent (Thermo Fisher Scientific) and purified by a silica column (Qiagen) to remove any excess DSS. One microgram of RNA was reverse transcribed to cDNA with SuperScript III First-Strand Synthesis System (Life Technologies). Transcript levels were analyzed using Power SYBR Green PCR Master Mix (ABI, 4367659) on a Step One Plus Real-Time PCR System (Life Technologies). Gene expression levels were quantitatively normalized to the expression of *Gapdh* by the change-in-cycling-threshold (ΔΔCT) method.

### Recombinant protein purification

The Retriever complex was expressed from *Sf9* cells using the Bac-to-Bac system (Thermo Fisher) and purified through Ni-NTA affinity, cation exchange, anion exchange, and size exclusion chromatographical steps, essentially as previously described. ^14^

GST- and MBP-tagged LRMDA and Rab proteins were expressed in *E. coli* BL21 (DE3) Gold cells (Sigma) cultured in Luria Broth. For MBP-tagged constructs, media were supplemented with 0.2% (w/v) glucose. For Rab proteins, media were supplemented with 2 mM MgCl₂. Cultures were grown at 37 °C to an OD₆₀₀ of 0.5, then induced with 1 mM IPTG (or 0.5 mM for different GST-Rab proteins), followed by overnight expression at 18 °C. Cells were harvested by centrifugation at 19,000 rpm for 30 min at 4 °C. Pellets were resuspended in 30 mL of lysis buffer [200 mM NaCl, 20 mM Tris-HCl (pH 8.0), 5 mM 2-mercaptoethanol (BME), 1 μg/mL antipain, 1 μg/mL leupeptin, 1 mM benzamidine, and 10% (w/v) glycerol]. Cells were lysed by five cycles of sonication on ice. GST-tagged proteins were purified using Glutathione Sepharose beads (Cytiva) and eluted with 100 mM Tris-HCl (pH 8.5) containing 30 mM reduced glutathione. MBP-tagged proteins were purified using amylose resin (New England Biolabs) and eluted with 20 mM Tris-HCl (pH 8.0), 5 mM BME, and 2% (w/v) maltose. Affinity-purified proteins were also purified over anion exchange chromatography on a 2-mL Source 15Q column (Cytiva) equilibrated with 20 mM Tris-HCl (pH 8.0) and 5 mM BME. Proteins were eluted with a 0–400 mM NaCl linear gradient over 40 mL. Final polishing was performed by size exclusion chromatography using a 24-mL Superdex Increase 200 column equilibrated with 10 mM HEPES (pH 7.0), 100 mM NaCl, 5% (w/v) glycerol, and 1 mM DTT. For Rab proteins, 2 mM MgCl₂ was included all purification buffers.

All chromatography steps were performed using Cytiva columns on an ÄKTA^TM^ Pure protein purification system. Sequences of recombinant proteins and DNA oligos are available in **Supplemental Table 3, 4**.

### GST pull-down assays

GST pull-down assays were performed essentially as previously described.^32^ Briefly, bait (100-200 pmol of GST-tagged proteins) and prey (100-1000 pmol for Retriever) were mixed with 20 µL of Glutathione Sepharose beads (Cytiva) in 1 mL of binding buffer (50 mM NaCl, 30 mM Tris pH 7.5, 5%(w/v) Glycerol, 5 mM MgCl_2_, 2 mM DTT, and 0.05% Triton X-100) at 4 °C for 30 min. After three 1-mL washes with the binding buffer, bound proteins were eluted with 100 mM Tris pH 8.5, 50 mM NaCl, and 30 mM reduced glutathione and examined by SDS-PAGE and Coomassie blue staining.

### GTPase nucleotide loading

Various Rab proteins were loaded with nucleotides (GMPPNP or GDP) before use, as previously described.^53^ Briefly, Rab proteins were mixed with 5 mM EDTA and 2 mM GMPPNP (or GDP). After 1 hr of incubation at 37 °C, 20 mM MgCl₂ was added to quench the reaction. The mixture was then stored on ice till use in pull-down assays.

### AlphaFold3 prediction and analysis

AlphaFold3 predictions are performed using the AlphaFold server (https://alphafoldserver.com).^25^ Typically, 25 models are obtained using random seeds and aligned in Pymol to examine the consistency. Reliability of the predicted models is assessed by the predicted local difference distance test (pLDDT) scores, the predicted aligned error (PAE) scores, and the manual examination of the consistency of 25 solutions.

### Listeria invasion into dendritic cells

DC2.4 cells were seeded at a density of 3 × 10⁵ cells per well in 8-well chamber slides one day prior to infection. On the day of infection, cells were washed with RPMI 1640 medium supplemented with 10% FBS but without antibiotics. The medium was then replaced with fresh antibiotic-free complete growth medium. *Listeria monocytogenes* (Microbiologics, Cat# 0687P) was washed to remove the bacterial broth and resuspended in RPMI medium without antibiotics. A total of 3 × 10⁶ PFU of *Listeria monocytogenes* was added to each well. Infected cells were incubated for 1.5 hours. After incubation, extracellular bacteria were removed by washing first with antibiotic-free medium and then with PBS. Cells were fixed with 4% paraformaldehyde (PFA) for 15 minutes and subsequently washed twice with PBS. Nuclei were stained with DAPI. Confocal images were acquired using a DMi8 microscope system (Leica Microsystems) equipped with a 63× oil immersion objective, with each image captured at a resolution of at least 2048 × 2048 pixels. Multichannel fluorescence imaging (four-color) was performed with well-separated channels for DAPI and EGFP. Each channel was sequentially scanned to minimize fluorophore crosstalk. Colocalization analysis was performed using Imaris image analysis software to quantify *Listeria monocytogenes* per cell.

### Histology and immunostaining

Freshly isolated colons were Swiss-rolled, fixed in formalin, and embedded in paraffin. H&E staining was conducted using a standard protocol by the UT Southwestern Histology core.

### Lamina propria leukocyte isolation

Lymphocytes from colonial lamina propria were isolated as previously described with minor changes.^22, 54^ Briefly, colons were flushed with the ice-cold PBS with 2% FBS, opened longitudinally, and cut into 1 cm pieces. The tissues were incubated with 15 ml of PBS with 1 mM EDTA, 1 mM DTT, and 2% FBS at 37°C in a shaker at 250 rpm for 15 min. The tissues were washed in PBS with 2% FBS and placed in RPMI-1640 with 2% FBS, 50 μg/ml of DNase I, and 50 μg/ml of Liberase. After incubation at 37°C for 30 min in a shaker at 250 rpm, the solution was vortexed and passed through a cell strainer. The cell suspensions were washed and applied to a 40%:80% Percoll gradient. After centrifugation at 800g for 20 min, lymphocytes were collected from the gradient interface.

### Single cell expression data

The gene expression data presented here was accessed from the Broad Single Cell Portal and analyzed data from ulcerative colitis as well as Crohn’s disease patients ^27, 28, 55^.

### Flow cytometry

Peripheral blood cells were isolated and red blood cell (RBC) lysis buffer was added to remove RBCs. Peripheral blood cells were washed with FACS staining buffer (PBS with 1% (w/v) BSA) and then centrifuged at 500 × g for 5 minutes. Peripheral blood cells were stained for 1 hour at 4°C, in 100 μl of a 1:200 cocktail of fluorescence-conjugated antibodies to 8 cell surface markers encompassing the major immune lineages: B220, CD19, CD3ε, CD4, CD8α, CD11b, F4/80, NK 1.1 and 1:200 Fc block. Lamina propria lymphocytes were stained at a 1:200 dilution in the presence of anti-mouseCD16/32 antibody for 30 min at 4°C with fluorochrome-conjugated antibodies against CD45, CD19, TCRβ, CD4, CD8α, MHC-II, CD11c and CD64. To analyze IL-17 production, lamina propria lymphocytes were stimulated with the Leukocyte Activation Cocktail, with BDGolgiPlug™ (BD) for 3 hours. For intracellular staining, cells were stained with surface markers and then fixed, permeabilized, and stained with antibodies specific for IL-17 or FOXP3.

## Data reproducibility and statistical analysis

All strains were generated and maintained on the same pure inbred background (C57BL/6J); experimental assessment of variance was not performed. No data was excluded. The investigator was not blinded to genotypes or group allocations during any experiment. Comparisons of differences were between two unpaired experimental groups in all cases. An unpaired *t*-test (Student’s *t* test) is appropriate and was used for such comparisons. One-way or two-way ANOVA with post hoc Tukey test was applied to experiments with three or more groups. The phenotype of mice (C57BL/6J) and primary cells of these mice are expected to follow a normal distribution. The statistical significance of differences between experimental groups was determined with GraphPad Prism 10 software and the Student’s *t* test (unpaired, two-tailed). A *P* value of less than 0.05 was considered statistically significant. No pre-specified effect size was assumed, and in general four mice or more for each genotype or condition were used in experiments; this sample size was sufficient to demonstrate statistically significant differences in comparisons between two unpaired experimental groups by an unpaired *t*-test.

## Acknowledgments

We thank the Proteomics core at UT Southwestern. GST-Rab constructs were a gift from Yongqun Zhu at Zhejiang University, China. Research was supported by funding from the National Institutes of Health (R35 GM128786 to B.C., R01 DK107733 to E.B., R01 HL179813 to B.C. and E.B., and R01 DK133229, R01 DK119360 to E.E.T.), and the National Science Foundation (CDF 2047640 to B.C.).

## Author Contributions

Conceptualization, E.B., B.C., B.B., and E.E.T.; Methodology, E.E.T., R.S., H.R., and C.N.; Investigation, R.S., C.N., A.S., D.A.K, A.D., D.J.B., Q.L., J.J.M., H.R.; Visualization, R.S. and B.C. Writing—original draft, E.E.T., E.B., R.S., and B.C..; Writing—review & editing, E.E.T., E.B., B.C. and R.S.; Funding acquisition, E.E.T., E.B. and B. C.; Supervision-E.E.T., B.C. and E.B.

## Material Availability

Further information and requests for resources and reagents should be directed to and will be fulfilled by the corresponding Contacts, E.E.T., E.B. or B.C.

---

**Extended Data Fig. 1. Generation and validation of *Lrmda*-deficient mice.**

**(a)** Gene targeting strategy for mouse *Lrmda*. Genomic sequences of the *Lrmda^+/+^* and *Lrmda^-/-^* alleles are shown. The *Lrmda^-/-^* allele was generated by CRISPR/Cas9-mediated deletion of 2 base pairs in exon 2. **(b)** Immunoblot analysis of bone marrow-derived dendritic cells (BMDCs) and bone marrow-derived macrophages (BMDMs) from *Lrmda^+/+^* and *Lrmda^-/-^* mice, using antibodies against VPS35L, CCDC93, COMMD1, and LRMDA. β-actin was used as a loading control.

**Extended Data Fig. 2. Expression of LRMDA in samples from IBD patients.**

**(a)** UMAP of single-cell RNA-seq data showing *Lrmda* expression in colon biopsies from Crohn’s disease patients. **(b, c)** *Lrmda* expression in small intestine (b) and colon (c) from IBD patients as determined by meta-analysis of RNA-seq datasets.^29^ Data are expressed as means ± s.d. and significance was determined by one-way ANOVA with multiple comparisons (ns, not significant, \*\*\*\**P*<0.0001).

**Extended Data Fig. 3. Immune homeostasis in *Lrmda*-deficient mice.**

**(a)** Flow cytometry analysis of peripheral blood from *Lrmda^+/+^* and *Lrmda^-/-^* mice, showing frequencies of B cells (CD19⁺ B220⁺), T cells (CD3⁺), CD4⁺ T cells, CD8⁺ T cells, neutrophils (CD11b⁺ F4/80⁻), macrophages (F4/80⁺), and NK cells (NK1.1⁺ CD3⁻). **(b-d)** Analysis of conventional dendritic cell (cDC) subsets in spleens from *Lrmda^+/+^* and *Lrmda^-/-^* mice. (b) Representative flow cytometry plots showing gating for cDC1 and cDC2 subsets. (c, d) Quantification of splenic cDC1 (c) and cDC2 (d) populations. **(e-h)** Analysis of plasmacytoid dendritic cells (pDCs) in spleens from *Lrmda^+/+^* and *Lrmda^-/-^* mice. (e) Representative flow cytometry plots showing the gating strategy for the pDC population. (f) Quantification of pDC frequencies in the spleen. (g) IFNα levels measured in culture supernatants of splenic pDCs following 24-hour stimulation with CpG. (h) Serum IFNα levels measured 6 hours after in vivo CpG injection. **(i)** Flow cytometry analysis of immune cell subsets in the colonic lamina propria from *Lrmda^+/+^* and *Lrmda^-/-^* mice, including B cells (CD19+), T cells (TCRb+), CD4+ T cells (TCRb+ CD4+), CD8+ T cells (TCRb+ CD8+), macrophages (CD64+ MHCII+), dendritic cells (CD11c+ MHCII+ CD64-). n=5 independent mice per genotype. **(j)** FOXP3 expression in colonic CD4+ T cells (TCRb+ CD4+). **(k)** IL-17A expression in colonic CD4⁺ T cells. Each data point represents an individual mouse of the indicated genotype. Data are expressed as means ± s.d. and significance was determined by unpaired Student t-test (a, c, d, f, g, h, i, j, k). ns, not significant. Data are representative of at least three independent experiments.

