## Supplemental Table 2 for "The Rab32-LRMDA-Retriever Complex is a Key Regulator of Intestinal Immune Homeostasis"

| **Name** | **Description** | **Source/Reference** | **Identifier** |
| --- | --- | --- | --- |
| **Recombinant protein preparation** | | | |
| pGexTev empty vector | pGexTev empty vector for bacterial expression of GST-Tev-tagged proteins | Published [PMID: [19363480](https://pubmed.ncbi.nlm.nih.gov/19363480/)] | pGexTev |
| pMalC2Tev empty vector | pMalC2Tev empty vector for bacterial expression of MBP-Tev-tagged proteins | Published [PMID: [19363480](https://pubmed.ncbi.nlm.nih.gov/19363480/)] | pMalTev |
| GST-hLRMDA | GST-Tev-(GGS_2_)-hLRMDA (1-226, full length, codon optimized for bacterial expression) in pGexTev vector | This Study | pCN36 |
| GST-hLRMDA point mutants | T152D  S174D  V169P/L203P  V169D/L203D  G204E/C206Y  R219G  R222* (deletion of D223, D224, Q225, and L226)  L226G  (all in pGexTev vector) | This study  This study  This study  This study  This study  This study  This study  This study | pDK458  pDK459  pDK461  pDK410  pCN53  pCN54  pCN113  pCN112 |
| GST-hLRMDA  GGSG linker mutants | V168/V169/K170/P171 GGSG; Q200/G201/V202/L203 GGSG, in pGexTev vector | This study | pDK462 |
| GST-hLRMDA NT | GST-Tev-(GGS_2_)-hLRMDA NT (1-167) in pGexTev vector | This study | pCN75 |
| GST-hLRMDA CT | GST-Tev-(GGS_2_)-hLRMDA CT (150-226) in pGexTev vector | This study | pCN76 |
| MBP-hLRMDA | MBP-Tev-(GGS_2_)-hLRMDA (1-226, full length, codon optimized for bacterial expression) in pMalC2Tev vector | This study | pCN77 |
| MBP-hLRMDA  Point mutants | D37A  N83K  K84E  Y106D  N117K  E155K  (all in pMalC2Tev vector) | This study  This study  This study  This study  This study  This study | pCN105  pCN61  pCN66  pCN67  pCN65  pCN62 |
| MBP-hLRMDA NT | MBP-Tev-(GGS_2_)-hLRMDA NT (1-167) in pMalC2Tev vector | This study | pCN44 |
| MBP-hLRMDA CT | MBP-Tev-(GGS_2_)-hLRMDA CT (150-226) in pMalC2Tev vector | This study | pCN45 |
| VPS35L | VPS35L (1-963, full-length, codon optimized for insect cell expression) in pAV5a vector | Published  [PMID: [39587083](https://pubmed.ncbi.nlm.nih.gov/39587083/)] | pCN38 |
| VPS26C | VPS26C (1-297, full-length) in pAV5a vector | Published [PMID: [34943955](https://pubmed.ncbi.nlm.nih.gov/34943955/)] | pCN39 |
| VPS29-His_6_ | VPS29-Tev-(GGS)_2_-His_6_ in pAV5a vector | Published [PMID: [34943955](https://pubmed.ncbi.nlm.nih.gov/34943955/)] | pCN40 |
| GST-hRab32 | GST-Tev-hRab32 (1-225, full length, codon optimized for bacterial expression) in pGexTev vector | This study | pCN43 |
| MBP-hRab32 | MBP-Tev-hRab32 (1-225, full length, codon optimized for bacterial expression) in pMalC2Tev vector | This study | pCN42 |
| GST-Rab5 | GST-TEV-hRab5A full length | Published  [PMID: 22939626] | pDK425 |
| GST-Rab11 | GST-thrombin-hRab11A full length | Published  [PMID: 22939626] | pDK431 |
| GST-Rab21 | GST-thrombin-hRab21 full length | Published  [PMID: 22939626] | pDK439 |
| GST-Rab38 | GST-thrombin-hRab38 full length | Published  [PMID: 22939626] | pDK453 |

Supplemental Table 2. DNA constructs used in this study.
