## Supplemental Table 3 for "The Rab32-LRMDA-Retriever Complex is a Key Regulator of Intestinal Immune Homeostasis"

Supplemental Table 3. Sequences of recombinant proteins used in this study

**GST-Tev-(GGS_2_)-hLRMDA FL(1-226) (added** WGGSGGS **to help Tev cleavage and Trp for concentration measurement)**

MSPILGYWKIKGLVQPTRLLLEYLEEKYEEHLYERDEGDKWRNKKFELGLEFPNLPYYIDGDVKLTQSMAIIRYIADKHNMLGGCPKERAEISMLEGAVLDIRYGVSRIAYSKDFETLKVDFLSKLPEMLKMFEDRLCHKTYLNGDHVTHPDFMLYDALDVVLYMDPMCLDAFPKLVCFKKRIEAIPQIDKYLKSSKYIAWPLQGWQATFGGGDHPPKSDLVPRGSENLYFQGHMWGGSGGSMAGLVVRGTQVSYIGQDCREIPEHLGRDCGHFAKRLDLSFNLLRSLEGLSAFRSLEELILDNNQLGDDLVLPGLPRLHTLTLNKNRITDLENLLDHLAEVTPALEYLSLLGNVACPNELVSLEKDEEDYKRYRCFVLYKLPNLKFLDAQKVTRQEREEALVRGVFMKVVKPKASSEDVASSPERHYTPLPSASRELTSHQGVLGKCRYVYYGKNSEGNRFIRDDQL

**GST-Tev-(GGS_2_)-hLRMDA^T152D^**

MSPILGYWKIKGLVQPTRLLLEYLEEKYEEHLYERDEGDKWRNKKFELGLEFPNLPYYIDGDVKLTQSMAIIRYIADKHNMLGGCPKERAEISMLEGAVLDIRYGVSRIAYSKDFETLKVDFLSKLPEMLKMFEDRLCHKTYLNGDHVTHPDFMLYDALDVVLYMDPMCLDAFPKLVCFKKRIEAIPQIDKYLKSSKYIAWPLQGWQATFGGGDHPPKSDLVPRGSENLYFQGHMWGGSGGSMAGLVVRGTQVSYIGQDCREIPEHLGRDCGHFAKRLDLSFNLLRSLEGLSAFRSLEELILDNNQLGDDLVLPGLPRLHTLTLNKNRITDLENLLDHLAEVTPALEYLSLLGNVACPNELVSLEKDEEDYKRYRCFVLYKLPNLKFLDAQKV**D**RQEREEALVRGVFMKVVKPKASSEDVASSPERHYTPLPSASRELTSHQGVLGKCRYVYYGKNSEGNRFIRDDQL

**GST-Tev-(GGS_2_)-hLRMDA^S174D^**

MSPILGYWKIKGLVQPTRLLLEYLEEKYEEHLYERDEGDKWRNKKFELGLEFPNLPYYIDGDVKLTQSMAIIRYIADKHNMLGGCPKERAEISMLEGAVLDIRYGVSRIAYSKDFETLKVDFLSKLPEMLKMFEDRLCHKTYLNGDHVTHPDFMLYDALDVVLYMDPMCLDAFPKLVCFKKRIEAIPQIDKYLKSSKYIAWPLQGWQATFGGGDHPPKSDLVPRGSENLYFQGHMWGGSGGSMAGLVVRGTQVSYIGQDCREIPEHLGRDCGHFAKRLDLSFNLLRSLEGLSAFRSLEELILDNNQLGDDLVLPGLPRLHTLTLNKNRITDLENLLDHLAEVTPALEYLSLLGNVACPNELVSLEKDEEDYKRYRCFVLYKLPNLKFLDAQKVTRQEREEALVRGVFMKVVKPKA**D**SEDVASSPERHYTPLPSASRELTSHQGVLGKCRYVYYGKNSEGNRFIRDDQL

**GST-Tev-(GGS_2_)-hLRMDA^V169P/L203P^**

MSPILGYWKIKGLVQPTRLLLEYLEEKYEEHLYERDEGDKWRNKKFELGLEFPNLPYYIDGDVKLTQSMAIIRYIADKHNMLGGCPKERAEISMLEGAVLDIRYGVSRIAYSKDFETLKVDFLSKLPEMLKMFEDRLCHKTYLNGDHVTHPDFMLYDALDVVLYMDPMCLDAFPKLVCFKKRIEAIPQIDKYLKSSKYIAWPLQGWQATFGGGDHPPKSDLVPRGSENLYFQGHMWGGSGGSMAGLVVRGTQVSYIGQDCREIPEHLGRDCGHFAKRLDLSFNLLRSLEGLSAFRSLEELILDNNQLGDDLVLPGLPRLHTLTLNKNRITDLENLLDHLAEVTPALEYLSLLGNVACPNELVSLEKDEEDYKRYRCFVLYKLPNLKFLDAQKVTRQEREEALVRGVFMKV**P**KPKASSEDVASSPERHYTPLPSASRELTSHQGV**P**GKCRYVYYGKNSEGNRFIRDDQL

**GST-Tev-(GGS_2_)-hLRMDA^V169D/L203D^**

MSPILGYWKIKGLVQPTRLLLEYLEEKYEEHLYERDEGDKWRNKKFELGLEFPNLPYYIDGDVKLTQSMAIIRYIADKHNMLGGCPKERAEISMLEGAVLDIRYGVSRIAYSKDFETLKVDFLSKLPEMLKMFEDRLCHKTYLNGDHVTHPDFMLYDALDVVLYMDPMCLDAFPKLVCFKKRIEAIPQIDKYLKSSKYIAWPLQGWQATFGGGDHPPKSDLVPRGSENLYFQGHMWGGSGGSMAGLVVRGTQVSYIGQDCREIPEHLGRDCGHFAKRLDLSFNLLRSLEGLSAFRSLEELILDNNQLGDDLVLPGLPRLHTLTLNKNRITDLENLLDHLAEVTPALEYLSLLGNVACPNELVSLEKDEEDYKRYRCFVLYKLPNLKFLDAQKVTRQEREEALVRGVFMKV**D**KPKASSEDVASSPERHYTPLPSASRELTSHQGV**D**GKCRYVYYGKNSEGNRFIRDDQL

**GST-Tev-(GGS_2_)-hLRMDA^G204E/C206Y^**

MSPILGYWKIKGLVQPTRLLLEYLEEKYEEHLYERDEGDKWRNKKFELGLEFPNLPYYIDGDVKLTQSMAIIRYIADKHNMLGGCPKERAEISMLEGAVLDIRYGVSRIAYSKDFETLKVDFLSKLPEMLKMFEDRLCHKTYLNGDHVTHPDFMLYDALDVVLYMDPMCLDAFPKLVCFKKRIEAIPQIDKYLKSSKYIAWPLQGWQATFGGGDHPPKSDLVPRGSENLYFQGHMWGGSGGSMAGLVVRGTQVSYIGQDCREIPEHLGRDCGHFAKRLDLSFNLLRSLEGLSAFRSLEELILDNNQLGDDLVLPGLPRLHTLTLNKNRITDLENLLDHLAEVTPALEYLSLLGNVACPNELVSLEKDEEDYKRYRCFVLYKLPNLKFLDAQKVTRQEREEALVRGVFMKVVKPKASSEDVASSPERHYTPLPSASRELTSHQGVL**E**K**Y**RYVYYGKNSEGNRFIRDDQL

**GST-Tev-(GGS_2_)-hLRMDA^R219G^**

MSPILGYWKIKGLVQPTRLLLEYLEEKYEEHLYERDEGDKWRNKKFELGLEFPNLPYYIDGDVKLTQSMAIIRYIADKHNMLGGCPKERAEISMLEGAVLDIRYGVSRIAYSKDFETLKVDFLSKLPEMLKMFEDRLCHKTYLNGDHVTHPDFMLYDALDVVLYMDPMCLDAFPKLVCFKKRIEAIPQIDKYLKSSKYIAWPLQGWQATFGGGDHPPKSDLVPRGSENLYFQGHMWGGSGGSMAGLVVRGTQVSYIGQDCREIPEHLGRDCGHFAKRLDLSFNLLRSLEGLSAFRSLEELILDNNQLGDDLVLPGLPRLHTLTLNKNRITDLENLLDHLAEVTPALEYLSLLGNVACPNELVSLEKDEEDYKRYRCFVLYKLPNLKFLDAQKVTRQEREEALVRGVFMKVVKPKASSEDVASSPERHYTPLPSASRELTSHQGVLGKCRYVYYGKNSEGN**G**FIRDDQL

**GST-Tev-(GGS_2_)-hLRMDA^R222*^**

MSPILGYWKIKGLVQPTRLLLEYLEEKYEEHLYERDEGDKWRNKKFELGLEFPNLPYYIDGDVKLTQSMAIIRYIADKHNMLGGCPKERAEISMLEGAVLDIRYGVSRIAYSKDFETLKVDFLSKLPEMLKMFEDRLCHKTYLNGDHVTHPDFMLYDALDVVLYMDPMCLDAFPKLVCFKKRIEAIPQIDKYLKSSKYIAWPLQGWQATFGGGDHPPKSDLVPRGSENLYFQGHMWGGSGGSMAGLVVRGTQVSYIGQDCREIPEHLGRDCGHFAKRLDLSFNLLRSLEGLSAFRSLEELILDNNQLGDDLVLPGLPRLHTLTLNKNRITDLENLLDHLAEVTPALEYLSLLGNVACPNELVSLEKDEEDYKRYRCFVLYKLPNLKFLDAQKVTRQEREEALVRGVFMKVVKPKASSEDVASSPERHYTPLPSASRELTSHQGVLGKCRYVYYGKNSEGNRFIR*****

**GST-Tev-(GGS_2_)-hLRMDA^VVKP->GGSG, QGVL->GGSG^**

MSPILGYWKIKGLVQPTRLLLEYLEEKYEEHLYERDEGDKWRNKKFELGLEFPNLPYYIDGDVKLTQSMAIIRYIADKHNMLGGCPKERAEISMLEGAVLDIRYGVSRIAYSKDFETLKVDFLSKLPEMLKMFEDRLCHKTYLNGDHVTHPDFMLYDALDVVLYMDPMCLDAFPKLVCFKKRIEAIPQIDKYLKSSKYIAWPLQGWQATFGGGDHPPKSDLVPRGSENLYFQGHMWGGSGGSMAGLVVRGTQVSYIGQDCREIPEHLGRDCGHFAKRLDLSFNLLRSLEGLSAFRSLEELILDNNQLGDDLVLPGLPRLHTLTLNKNRITDLENLLDHLAEVTPALEYLSLLGNVACPNELVSLEKDEEDYKRYRCFVLYKLPNLKFLDAQKVTRQEREEALVRGVFMK**GGSG**KASSEDVASSPERHYTPLPSASRELTSH**GGSG**GKCRYVYYGKNSEGNRFIRDDQL

**GST-Tev-(GGS_2_)-hLRMDA^L226G^**

MSPILGYWKIKGLVQPTRLLLEYLEEKYEEHLYERDEGDKWRNKKFELGLEFPNLPYYIDGDVKLTQSMAIIRYIADKHNMLGGCPKERAEISMLEGAVLDIRYGVSRIAYSKDFETLKVDFLSKLPEMLKMFEDRLCHKTYLNGDHVTHPDFMLYDALDVVLYMDPMCLDAFPKLVCFKKRIEAIPQIDKYLKSSKYIAWPLQGWQATFGGGDHPPKSDLVPRGSENLYFQGHMWGGSGGSMAGLVVRGTQVSYIGQDCREIPEHLGRDCGHFAKRLDLSFNLLRSLEGLSAFRSLEELILDNNQLGDDLVLPGLPRLHTLTLNKNRITDLENLLDHLAEVTPALEYLSLLGNVACPNELVSLEKDEEDYKRYRCFVLYKLPNLKFLDAQKVTRQEREEALVRGVFMKVVKPKASSEDVASSPERHYTPLPSASRELTSHQGVLGKCRYVYYGKNSEGNRFIRDDQ**G**

**GST-Tev-(GGS_2_)-hLRMDA NT(1-167)**

MSPILGYWKIKGLVQPTRLLLEYLEEKYEEHLYERDEGDKWRNKKFELGLEFPNLPYYIDGDVKLTQSMAIIRYIADKHNMLGGCPKERAEISMLEGAVLDIRYGVSRIAYSKDFETLKVDFLSKLPEMLKMFEDRLCHKTYLNGDHVTHPDFMLYDALDVVLYMDPMCLDAFPKLVCFKKRIEAIPQIDKYLKSSKYIAWPLQGWQATFGGGDHPPKSDLVPRGSENLYFQGHMWGGSGGSMAGLVVRGTQVSYIGQDCREIPEHLGRDCGHFAKRLDLSFNLLRSLEGLSAFRSLEELILDNNQLGDDLVLPGLPRLHTLTLNKNRITDLENLLDHLAEVTPALEYLSLLGNVACPNELVSLEKDEEDYKRYRCFVLYKLPNLKFLDAQKVTRQEREEALVRGVFMK

**GST-Tev-(GGS_2_)-hLRMDA CT(150-226)**

MSPILGYWKIKGLVQPTRLLLEYLEEKYEEHLYERDEGDKWRNKKFELGLEFPNLPYYIDGDVKLTQSMAIIRYIADKHNMLGGCPKERAEISMLEGAVLDIRYGVSRIAYSKDFETLKVDFLSKLPEMLKMFEDRLCHKTYLNGDHVTHPDFMLYDALDVVLYMDPMCLDAFPKLVCFKKRIEAIPQIDKYLKSSKYIAWPLQGWQATFGGGDHPPKSDLVPRGSENLYFQGHMWGGSGGSKVTRQEREEALVRGVFMKVVKPKASSEDVASSPERHYTPLPSASRELTSHQGVLGKCRYVYYGKNSEGNRFIRDDQL

**MBP-Tev-(GGS_2_)-hLRMDA FL (1-226)**

MKTEEGKLVIWINGDKGYNGLAEVGKKFEKDTGIKVTVEHPDKLEEKFPQVAATGDGPDIIFWAHDRFGGYAQSGLLAEITPDKAFQDKLYPFTWDAVRYNGKLIAYPIAVEALSLIYNKDLLPNPPKTWEEIPALDKELKAKGKSALMFNLQEPYFTWPLIAADGGYAFKYENGKYDIKDVGVDNAGAKAGLTFLVDLIKNKHMNADTDYSIAEAAFNKGETAMTINGPWAWSNIDTSKVNYGVTVLPTFKGQPSKPFVGVLSAGINAASPNKELAKEFLENYLLTDEGLEAVNKDKPLGAVALKSYEEELAKDPRIAATMENAQKGEIMPNIPQMSAFWYAVRTAVINAASGRQTVDEALKDAQTNSSSNNNNNNNNNNLGIEGRISEFENLYFQGHWGGSGGSMAGLVVRGTQVSYIGQDCREIPEHLGRDCGHFAKRLDLSFNLLRSLEGLSAFRSLEELILDNNQLGDDLVLPGLPRLHTLTLNKNRITDLENLLDHLAEVTPALEYLSLLGNVACPNELVSLEKDEEDYKRYRCFVLYKLPNLKFLDAQKVTRQEREEALVRGVFMKVVKPKASSEDVASSPERHYTPLPSASRELTSHQGVLGKCRYVYYGKNSEGNRFIRDDQL

**MBP-Tev-(GGS_2_)-hLRMDA^D37A^**

MKTEEGKLVIWINGDKGYNGLAEVGKKFEKDTGIKVTVEHPDKLEEKFPQVAATGDGPDIIFWAHDRFGGYAQSGLLAEITPDKAFQDKLYPFTWDAVRYNGKLIAYPIAVEALSLIYNKDLLPNPPKTWEEIPALDKELKAKGKSALMFNLQEPYFTWPLIAADGGYAFKYENGKYDIKDVGVDNAGAKAGLTFLVDLIKNKHMNADTDYSIAEAAFNKGETAMTINGPWAWSNIDTSKVNYGVTVLPTFKGQPSKPFVGVLSAGINAASPNKELAKEFLENYLLTDEGLEAVNKDKPLGAVALKSYEEELAKDPRIAATMENAQKGEIMPNIPQMSAFWYAVRTAVINAASGRQTVDEALKDAQTNSSSNNNNNNNNNNLGIEGRISEFENLYFQGHWGGSGGSMAGLVVRGTQVSYIGQDCREIPEHLGRDCGHFAKRL**A**LSFNLLRSLEGLSAFRSLEELILDNNQLGDDLVLPGLPRLHTLTLNKNRITDLENLLDHLAEVTPALEYLSLLGNVACPNELVSLEKDEEDYKRYRCFVLYKLPNLKFLDAQKVTRQEREEALVRGVFMKVVKPKASSEDVASSPERHYTPLPSASRELTSHQGVLGKCRYVYYGKNSEGNRFIRDDQL

**MBP-Tev-(GGS_2_)-hLRMDA^N83K^**

MKTEEGKLVIWINGDKGYNGLAEVGKKFEKDTGIKVTVEHPDKLEEKFPQVAATGDGPDIIFWAHDRFGGYAQSGLLAEITPDKAFQDKLYPFTWDAVRYNGKLIAYPIAVEALSLIYNKDLLPNPPKTWEEIPALDKELKAKGKSALMFNLQEPYFTWPLIAADGGYAFKYENGKYDIKDVGVDNAGAKAGLTFLVDLIKNKHMNADTDYSIAEAAFNKGETAMTINGPWAWSNIDTSKVNYGVTVLPTFKGQPSKPFVGVLSAGINAASPNKELAKEFLENYLLTDEGLEAVNKDKPLGAVALKSYEEELAKDPRIAATMENAQKGEIMPNIPQMSAFWYAVRTAVINAASGRQTVDEALKDAQTNSSSNNNNNNNNNNLGIEGRISEFENLYFQGHWGGSGGSMAGLVVRGTQVSYIGQDCREIPEHLGRDCGHFAKRLDLSFNLLRSLEGLSAFRSLEELILDNNQLGDDLVLPGLPRLHTLTL**K**KNRITDLENLLDHLAEVTPALEYLSLLGNVACPNELVSLEKDEEDYKRYRCFVLYKLPNLKFLDAQKVTRQEREEALVRGVFMKVVKPKASSEDVASSPERHYTPLPSASRELTSHQGVLGKCRYVYYGKNSEGNRFIRDDQL

**MBP-Tev-(GGS_2_)-hLRMDA^K84E^**

MKTEEGKLVIWINGDKGYNGLAEVGKKFEKDTGIKVTVEHPDKLEEKFPQVAATGDGPDIIFWAHDRFGGYAQSGLLAEITPDKAFQDKLYPFTWDAVRYNGKLIAYPIAVEALSLIYNKDLLPNPPKTWEEIPALDKELKAKGKSALMFNLQEPYFTWPLIAADGGYAFKYENGKYDIKDVGVDNAGAKAGLTFLVDLIKNKHMNADTDYSIAEAAFNKGETAMTINGPWAWSNIDTSKVNYGVTVLPTFKGQPSKPFVGVLSAGINAASPNKELAKEFLENYLLTDEGLEAVNKDKPLGAVALKSYEEELAKDPRIAATMENAQKGEIMPNIPQMSAFWYAVRTAVINAASGRQTVDEALKDAQTNSSSNNNNNNNNNNLGIEGRISEFENLYFQGHWGGSGGSMAGLVVRGTQVSYIGQDCREIPEHLGRDCGHFAKRLDLSFNLLRSLEGLSAFRSLEELILDNNQLGDDLVLPGLPRLHTLTLN**E**NRITDLENLLDHLAEVTPALEYLSLLGNVACPNELVSLEKDEEDYKRYRCFVLYKLPNLKFLDAQKVTRQEREEALVRGVFMKVVKPKASSEDVASSPERHYTPLPSASRELTSHQGVLGKCRYVYYGKNSEGNRFIRDDQL

**MBP-Tev-(GGS_2_)-hLRMDA^Y106D^**

MKTEEGKLVIWINGDKGYNGLAEVGKKFEKDTGIKVTVEHPDKLEEKFPQVAATGDGPDIIFWAHDRFGGYAQSGLLAEITPDKAFQDKLYPFTWDAVRYNGKLIAYPIAVEALSLIYNKDLLPNPPKTWEEIPALDKELKAKGKSALMFNLQEPYFTWPLIAADGGYAFKYENGKYDIKDVGVDNAGAKAGLTFLVDLIKNKHMNADTDYSIAEAAFNKGETAMTINGPWAWSNIDTSKVNYGVTVLPTFKGQPSKPFVGVLSAGINAASPNKELAKEFLENYLLTDEGLEAVNKDKPLGAVALKSYEEELAKDPRIAATMENAQKGEIMPNIPQMSAFWYAVRTAVINAASGRQTVDEALKDAQTNSSSNNNNNNNNNNLGIEGRISEFENLYFQGHWGGSGGSMAGLVVRGTQVSYIGQDCREIPEHLGRDCGHFAKRLDLSFNLLRSLEGLSAFRSLEELILDNNQLGDDLVLPGLPRLHTLTLNKNRITDLENLLDHLAEVTPALE**D**LSLLGNVACPNELVSLEKDEEDYKRYRCFVLYKLPNLKFLDAQKVTRQEREEALVRGVFMKVVKPKASSEDVASSPERHYTPLPSASRELTSHQGVLGKCRYVYYGKNSEGNRFIRDDQL

**MBP-Tev-(GGS_2_)-hLRMDA^N117K^**

MKTEEGKLVIWINGDKGYNGLAEVGKKFEKDTGIKVTVEHPDKLEEKFPQVAATGDGPDIIFWAHDRFGGYAQSGLLAEITPDKAFQDKLYPFTWDAVRYNGKLIAYPIAVEALSLIYNKDLLPNPPKTWEEIPALDKELKAKGKSALMFNLQEPYFTWPLIAADGGYAFKYENGKYDIKDVGVDNAGAKAGLTFLVDLIKNKHMNADTDYSIAEAAFNKGETAMTINGPWAWSNIDTSKVNYGVTVLPTFKGQPSKPFVGVLSAGINAASPNKELAKEFLENYLLTDEGLEAVNKDKPLGAVALKSYEEELAKDPRIAATMENAQKGEIMPNIPQMSAFWYAVRTAVINAASGRQTVDEALKDAQTNSSSNNNNNNNNNNLGIEGRISEFENLYFQGHWGGSGGSMAGLVVRGTQVSYIGQDCREIPEHLGRDCGHFAKRLDLSFNLLRSLEGLSAFRSLEELILDNNQLGDDLVLPGLPRLHTLTLNKNRITDLENLLDHLAEVTPALEYLSLLGNVACP**K**ELVSLEKDEEDYKRYRCFVLYKLPNLKFLDAQKVTRQEREEALVRGVFMKVVKPKASSEDVASSPERHYTPLPSASRELTSHQGVLGKCRYVYYGKNSEGNRFIRDDQL

**MBP-Tev-(GGS_2_)-hLRMDA^E155K^**

MKTEEGKLVIWINGDKGYNGLAEVGKKFEKDTGIKVTVEHPDKLEEKFPQVAATGDGPDIIFWAHDRFGGYAQSGLLAEITPDKAFQDKLYPFTWDAVRYNGKLIAYPIAVEALSLIYNKDLLPNPPKTWEEIPALDKELKAKGKSALMFNLQEPYFTWPLIAADGGYAFKYENGKYDIKDVGVDNAGAKAGLTFLVDLIKNKHMNADTDYSIAEAAFNKGETAMTINGPWAWSNIDTSKVNYGVTVLPTFKGQPSKPFVGVLSAGINAASPNKELAKEFLENYLLTDEGLEAVNKDKPLGAVALKSYEEELAKDPRIAATMENAQKGEIMPNIPQMSAFWYAVRTAVINAASGRQTVDEALKDAQTNSSSNNNNNNNNNNLGIEGRISEFENLYFQGHWGGSGGSMAGLVVRGTQVSYIGQDCREIPEHLGRDCGHFAKRLDLSFNLLRSLEGLSAFRSLEELILDNNQLGDDLVLPGLPRLHTLTLNKNRITDLENLLDHLAEVTPALEYLSLLGNVACPNELVSLEKDEEDYKRYRCFVLYKLPNLKFLDAQKVTRQ**K**REEALVRGVFMKVVKPKASSEDVASSPERHYTPLPSASRELTSHQGVLGKCRYVYYGKNSEGNRFIRDDQL

**MBP-Tev-(GGS_2_)-hLRMDA NT(1-167)**

MKTEEGKLVIWINGDKGYNGLAEVGKKFEKDTGIKVTVEHPDKLEEKFPQVAATGDGPDIIFWAHDRFGGYAQSGLLAEITPDKAFQDKLYPFTWDAVRYNGKLIAYPIAVEALSLIYNKDLLPNPPKTWEEIPALDKELKAKGKSALMFNLQEPYFTWPLIAADGGYAFKYENGKYDIKDVGVDNAGAKAGLTFLVDLIKNKHMNADTDYSIAEAAFNKGETAMTINGPWAWSNIDTSKVNYGVTVLPTFKGQPSKPFVGVLSAGINAASPNKELAKEFLENYLLTDEGLEAVNKDKPLGAVALKSYEEELAKDPRIAATMENAQKGEIMPNIPQMSAFWYAVRTAVINAASGRQTVDEALKDAQTNSSSNNNNNNNNNNLGIEGRISEFENLYFQGHMWGGSGGSMAGLVVRGTQVSYIGQDCREIPEHLGRDCGHFAKRLDLSFNLLRSLEGLSAFRSLEELILDNNQLGDDLVLPGLPRLHTLTLNKNRITDLENLLDHLAEVTPALEYLSLLGNVACPNELVSLEKDEEDYKRYRCFVLYKLPNLKFLDAQKVTRQEREEALVRGVFMK

**MBP-Tev-(GGS_2_)-hLRMDA CT (150-226)**

MKTEEGKLVIWINGDKGYNGLAEVGKKFEKDTGIKVTVEHPDKLEEKFPQVAATGDGPDIIFWAHDRFGGYAQSGLLAEITPDKAFQDKLYPFTWDAVRYNGKLIAYPIAVEALSLIYNKDLLPNPPKTWEEIPALDKELKAKGKSALMFNLQEPYFTWPLIAADGGYAFKYENGKYDIKDVGVDNAGAKAGLTFLVDLIKNKHMNADTDYSIAEAAFNKGETAMTINGPWAWSNIDTSKVNYGVTVLPTFKGQPSKPFVGVLSAGINAASPNKELAKEFLENYLLTDEGLEAVNKDKPLGAVALKSYEEELAKDPRIAATMENAQKGEIMPNIPQMSAFWYAVRTAVINAASGRQTVDEALKDAQTNSSSNNNNNNNNNNLGIEGRISEFENLYFQGHMWGGSGGSKVTRQEREEALVRGVFMKVVKPKASSEDVASSPERHYTPLPSASRELTSHQGVLGKCRYVYYGKNSEGNRFIRDDQL

**VPS35L**

MAVFPWHSRNRNYKAEFASCRLEAVPLEFGDYHPLKPITVTESKTKKVNRKGSTSSTSSSSSSSVVDPLSSVLDGTDPLSMFAATADPAALAAAMDSSRRKRDRDDNSVVGSDFEPWTNKRGEILARYTTTEKLSINLFMGSEKGKAGTATLAMSEKVRTRLEELDDFEEGSQKELLNLTQQDYVNRIEELNQSLKDAWASDQKVKALKIVIQCSKLLSDTSVIQFYPSKFVLITDILDTFGKLVYERIFSMCVDSRSVLPDHFSPENANDTAKETCLNWFFKIASIRELIPRFYVEASILKCNKFLSKTGISECLPRLTCMIRGIGDPLVSVYARAYLCRVGMEVAPHLKETLNKNFFDFLLTFKQIHGDTVQNQLVVQGVELPSYLPLYPPAMDWIFQCISYHAPEALLTEMMERCKKLGNNALLLNSVMSAFRAEFIATRSMDFIGMIKECDESGFPKHLLFRSLGLNLALADPPESDRLQILNEAWKVITKLKNPQDYINCAEVWVEYTCKHFTKREVNTVLADVIKHMTPDRAFEDSYPQLQLIIKKVIAHFHDFSVLFSVEKFLPFLDMFQKESVRVEVCKCIMDAFIKHQQEPTKDPVILNALLHVCKTMHDSVNALTLEDEKRMLSYLINGFIKMVSFGRDFEQQLSFYVESRSMFCNLEPVLVQLIHSVNRLAMETRKVMKGNHSRKTAAFVRACVAYCFITIPSLAGIFTRLNLYLHSGQVALANQCLSQADAFFKAAISLVPEVPKMINIDGKMRPSESFLLEFLCNFFSTLLIVPDHPEHGVLFLVRELLNVIQDYTWEDNSDEKIRIYTCVLHLLSAMSQETYLYHIDKVDSNDSLYGGDSKFLAENNKLCETVMAQILEHLKTLAKDEALKRQSSLGLSFFNSILAHGDLRNNKLNQLSVNLWHLAQRHGCADTRTMVKTLEYIKKQSKQPDMTHLTELALRLPLQTRT

**VPS26C**

MGTALDIKIKRANKVYHAGEVLSGVVVISSKDSVQHQGVSLTMEGTVNLQLSAKSVGVFEAFYNSVKPIQIINSTIEMVKPGKFPSGKTEIPFEFPLHLKGNKVLYETYHGVFVNIQYTLRCDMKRSLLAKDLTKTCEFIVHSAPQKGKFTPSPVDFTITPETLQNVKERALLPKFLLRGHLNSTNCVITQPLTGELVVESSEAAIRSVELQLVRVETCGCAEGYARDATEIQNIQIADGDVCRGLSVPIYMVFPRLFTCPTLETTNFKVEFEVNIVVLLHPDHLITENFPLKLCRI

**VPS29-Tev-(GGS_2_)-His_6_** MAGHRLVLVLGDLHIPHRCNSLPAKFKKLLVPGKIQHILCTGNLCTKESYDYLKTLAGDVHIVRGDFDENLNYPEQKVVTVGQFKIGLIHGHQVIPWGDMASLALLQRQFDVDILISGHTHKFEAFEHENKFYINPGSATGAYNALETNIIPSFVLMDIQASTVVTYVYQLIGDDVKVERIEYKKPENLYFQGGGSGGSHHHHHH

**GST-Tev-hRab32 FL (1-225)**

MSPILGYWKIKGLVQPTRLLLEYLEEKYEEHLYERDEGDKWRNKKFELGLEFPNLPYYIDGDVKLTQSMAIIRYIADKHNMLGGCPKERAEISMLEGAVLDIRYGVSRIAYSKDFETLKVDFLSKLPEMLKMFEDRLCHKTYLNGDHVTHPDFMLYDALDVVLYMDPMCLDAFPKLVCFKKRIEAIPQIDKYLKSSKYIAWPLQGWQATFGGGDHPPKSDLVPRGSENLYFQGHMAGGGAGDPGLGAAAAPAPETREHLFKVLVIGELGVGKTSIIKRYVHQLFSQHYRATIGVDFALKVLNWDSRTLVRLQLWDIAGQERFGNMTRVYYKEAVGAFVVFDISRSSTFEAVLKWKSDLDSKVHLPNGSPIPAVLLANKCDQNKDSSQSPSQVDQFCKEHGFAGWFETSAKDNINIEEAARFLVEKILVNHQSFPNEENDVDKIKLDQETLRAENKSQCC

**MBP-Tev-hRab32 FL (1-225)**

MKTEEGKLVIWINGDKGYNGLAEVGKKFEKDTGIKVTVEHPDKLEEKFPQVAATGDGPDIIFWAHDRFGGYAQSGLLAEITPDKAFQDKLYPFTWDAVRYNGKLIAYPIAVEALSLIYNKDLLPNPPKTWEEIPALDKELKAKGKSALMFNLQEPYFTWPLIAADGGYAFKYENGKYDIKDVGVDNAGAKAGLTFLVDLIKNKHMNADTDYSIAEAAFNKGETAMTINGPWAWSNIDTSKVNYGVTVLPTFKGQPSKPFVGVLSAGINAASPNKELAKEFLENYLLTDEGLEAVNKDKPLGAVALKSYEEELAKDPRIAATMENAQKGEIMPNIPQMSAFWYAVRTAVINAASGRQTVDEALKDAQTNSSSNNNNNNNNNNLGIEGRISEFENLYFQGHMAGGGAGDPGLGAAAAPAPETREHLFKVLVIGELGVGKTSIIKRYVHQLFSQHYRATIGVDFALKVLNWDSRTLVRLQLWDIAGQERFGNMTRVYYKEAVGAFVVFDISRSSTFEAVLKWKSDLDSKVHLPNGSPIPAVLLANKCDQNKDSSQSPSQVDQFCKEHGFAGWFETSAKDNINIEEAARFLVEKILVNHQSFPNEENDVDKIKLDQETLRAENKSQCC

**GST-Tev-Rab5a**

MSPILGYWKIKGLVQPTRLLLEYLEEKYEEHLYERDEGDKWRNKKFELGLEFPNLPYYIDGDVKLTQSMAIIRYIADKHNMLGGCPKERAEISMLEGAVLDIRYGVSRIAYSKDFETLKVDFLSKLPEMLKMFEDRLCHKTYLNGDHVTHPDFMLYDALDVVLYMDPMCLDAFPKLVCFKKRIEAIPQIDKYLKSSKYIAWPLQGWQATFGGGDHPPKSDLVPRGSENLYFQGMASRGATRPNGPNTGNKICQFKLVLLGESAVGKSSLVLRFVKGQFHEFQESTIGAAFLTQTVCLDDTTVKFEIWDTAGQERYHSLAPMYYRGAQAAIVVYDITNEESFARAKNWVKELQRQASPNIVIALSGNKADLANKRAVDFQEAQSYADDNSLLFMETSAKTSMNVNEIFMAIAKKLPKNEPQNPGANSARGRGVDLTEPTQPTRNQCCSN

**GST-Thrombin-Rab11a**

MSPILGYWKIKGLVQPTRLLLEYLEEKYEEHLYERDEGDKWRNKKFELGLEFPNLPYYIDGDVKLTQSMAIIRYIADKHNMLGGCPKERAEISMLEGAVLDIRYGVSRIAYSKDFETLKVDFLSKLPEMLKMFEDRLCHKTYLNGDHVTHPDFMLYDALDVVLYMDPMCLDAFPKLVCFKKRIEAIPQIDKYLKSSKYIAWPLQGWQATFGGGDHPPKSDLVPRGSGTRDDEYDYLFKVVLIGDSGVGKSNLLSRFTRNEFNLESKSTIGVEFATRSIQVDGKTIKAQIWDTAGQERYRAITSAYYRGAVGALLVYDIAKHLTYENVERWLKELRDHADSNIVIMLVGNKSDLRHLRAVPTDEARAFAEKNGLSFIETSALDSTNVEAAFQTILTEIYRIVSQKQMSDRRENDMSPSNNVVPIHVPPTTENKPKVQCCQNI

**GST-Thrombin-Rab21**

MSPILGYWKIKGLVQPTRLLLEYLEEKYEEHLYERDEGDKWRNKKFELGLEFPNLPYYIDGDVKLTQSMAIIRYIADKPNMLGGCPKERAEISMLEGAVLDIRYGVSRIAYSKDFETLKVDFLSKLPEMLKMFEDRLCHKTYLNGDHVTHPDFMLYDALDVVLYMDPMCLDAFPKLVCFKKRIEAIPQIDKYLKSSKYIAWPLQGWQATFGGGDHPPKSDLVPRGSRAYSFKVVLLGEGCVGKTSLVLRYCENKFNDKHITTLQASFLTKKLNIGGKRVNLAIWDTAGQERFHALGPIYYRDSNGAILVYDITDEDSFQKVKNWVKELRKMLGNEICLCIVGNKIDLEKERHVSIQEAESYAESVGAKHYHTSAKQNKGIEELFLDLCKRMIETAQVDERAKGNGSSQPGTARRGVQIIDDEPQAQTSGGGCCSSG

**GST-Thrombin-Rab38**

MSPILGYWKIKGLVQPTRLLLEYLEEKYEEHLYERDEGDKWRNKKFELGLEFPNLPYYIDGDVKLTQSMAIIRYIADKHNMLGGCPKERAEISMLEGAVLDIRYGVSRIAYSKDFETLKVDFLSKLPEMLKMFEDRLCHKTYLNGDHVTHPDFMLYDALDVVLYMDPMCLDAFPKLVCFKKRIEAIPQIDKYLKSSKYIAWPLQGWQATFGGGDHPPKSDLVPRGSQAPHKEHLYKLLVIGDLGVGKTSIIKRYVHQNFSSHYRATIGVDFALKVLHWDPETVVRLQLWDIAGQERFGNMTRVYYREAMGAFIVFDVTRPATFEAVAKWKNDLDSKLSLPNGKPVSVVLLANKCDQGKDVLMNNGLKMDQFCKEHGFVGWFETSAKENINIDEASRCLVKHILANECDLMESIEPDVVKPHLTSTKVASCSGCAKS
