## Supplemental Table 4 for "The Rab32-LRMDA-Retriever Complex is a Key Regulator of Intestinal Immune Homeostasis"

Supplemental Table 4. DNA oligos used in this study

| GST-Tev-(GGS_2_)-hLRMDA | Ordered from ThermoFisher GeneArt |
| --- | --- |
| GST-Tev-(GGS_2_)-hLRMDA **^T152D^** | Cbyo-230926-5, GACCGTCAAGAACGTGAAGAGGCAC, Aliblunt for LRMDAopti T152D fw  CTGAAATTTCTGGATGCCCAGAAAGTT  Cbyo-230926-6, AACTTTCTGGGCATCCAGAAATTTCAG, Aliblunt for LRMDAopti T152D bw |
| GST-Tev-(GGS_2_)-hLRMDA **^S174D^** | Cbyo-230926-7, GACAGCGAAGATGTTGCAAGCAG, Aliblunt for LRMDAopti S174D fw  GTTTTTATGAAAGTTGTTAAACCGAAAGCC  Cbyo-230926-8, GGCTTTCGGTTTAACAACTTTCATAAAAAC, Aliblunt for LRMDAopti S174D bw |
| GST-Tev-(GGS_2_)-hLRMDA **^V169P/L203P^** | Ordered from ThermoFisher GeneArt |
| GST-Tev-(GGS_2_)-hLRMDA **^V169D/L203D^** | Ordered from ThermoFisher GeneArt |
| GST-Tev-(GGS_2_)-hLRMDA **^G204E/C206Y^** | CNo240908-5, ATATCGTTATGTGTATTATGGCAAAAACAGC, Fw primer to make G204E/C206Y  CAGCCATCAGGGTGTTCTGGAAAA  CNo240908-6, TTTTCCAGAACACCCTGATGGCTG, Bw primer to make G204E/C206Y |
| GST-Tev-(GGS_2_)-hLRMDA **^R219G^** | CNo240908-7, GGTTTTATTCGTGATGATCAGCTGTAAGG, Fw primer to make R219G  GCAAAAACAGCGAAGGCAAC  CN0240908-8, GTTGCCTTCGCTGTTTTTGC, Bw primer to make R219G |
| GST-Tev-(GGS_2_)-hLRMDA **^R222*^** | CNo250214-11fw, TAAGGATCCGCGGCCGCATC, Fw primer to make R222*  CAGCGAAGGCAACCGTTTTATTCGT  CNo250214-12bw, ACGAATAAAACGGTTGCCTTCGCTG, Bw primer to make R222* |
| GST-Tev-(GGS_2_)-hLRMDA **^L226G^** | Use CNo250214-11fw, as fw primer to make L226G  GAAGGCAACCGTTTTATTCGTGATGATCAGGGC  CNo250214-11bw, GCCCTGATCATCACGAATAAAACGGTTGCCTTC, as bw primer to make L226G |
| GST-Tev-(GGS_2_)-hLRMDA ^VVKP->GGSG/QGVL->GGSG^ | Ordered from ThermoFisher GeneArt |
| GST-Tev-(GGS_2_)-hLRMDA NT (1-167) | Cbyo-230926-1, TAAGGATCCGCGGCCGCATC, Aliblunt for LRMDAopti 1-167 fw  GCACTGGTTCGTGGTGTTTTTATGAAA  Cbyo-230926-2, TTTCATAAAAACACCACGAACCAGTGC, Aliblunt for LRMDAopti 1-167 bw |
| GST-Tev-(GGS_2_)-hLRMDA CT (150-226) | Cbyo-230926-3, AAAGTTACCCGTCAAGAACGTGAAG, Aliblunt for LRMDAopti 150-226 fw  gTGGGGTGGTAGCGGTGGTAGT  Cbyo-230926-4, ACTACCACCGCTACCACCCCAc, Aliblunt for LRMDAopti 150-226 bw |
| MBP-Tev-(GGS_2_)-hLRMDA **^D37A^** | CNo250214-1, CACTGTCATTTAATCTGCTGCGTAGC, Aliblunt Fw primer to make MBP-LRMDA D37A,  *GTGGTCATTTTGCCAAACGTCTGG*  CNo250214-2, CCAGACGTTTGGCAAAATGACCAC, Aliblunt Bw primer to make MBP-LRMDA D37A |
| MBP-Tev-(GGS_2_)-hLRMDA **^N83K^** | CNo240908-1, GAAAAACCGTATTACCGATCTGGAAAAC, FW PRIMER TO MAKE N83K  GTCTGCATACCCTGACACTGAA  CN0240908-2, TTCAGTGTCAGGGTATGCAGAC, Bw primer to make N83K |
| MBP-Tev-(GGS_2_)-hLRMDA **^K84E^** | CNo240908-9, GAAAACCGTATTACCGATCTGGAAAACC, Fw primer to make K84E  GTCTGCATACCCTGACACTGAAT  CNo240908-10, ATTCAGTGTCAGGGTATGCAGAC, Bw primer to make K84E |
| MBP-Tev-(GGS_2_)-hLRMDA **^Y106D^** | CNo240908-11, GATCTGAGCCTGCTGGGTAATG, Fw primer to make Y106D  GAAGTTACACCGGCACTGGAA  CNo240908-12, TTCCAGTGCCGGTGTAACTTC, Bw primer to make Y106D |
| MBP-Tev-(GGS_2_)-hLRMDA **^N117K^** | CNo240920-1, GGAACTGGTGAGCCTGGAAAAAG, Fw primer to make N117K  CTGGGTAATGTTGCATGTCCGAA  CNo240920-2, TTCGGACATGCAACATTACCCAG, Bw primer to make N117K |
| MBP-Tev-(GGS_2_)-hLRMDA **^E155K^** | CNo240908-3, AAACGTGAAGAGGCACTGGTTC, Fw primer to make E155K  GATGCCCAGAAAGTTACCCGTCAA  CNo240908-4, TTGACGGGTAACTTTCTGGGCATC, Bw primer to make E155K |
