## Supplementary figures and images for "The Rab32-LRMDA-Retriever Complex is a Key Regulator of Intestinal Immune Homeostasis"

### Extended Data Figure 1

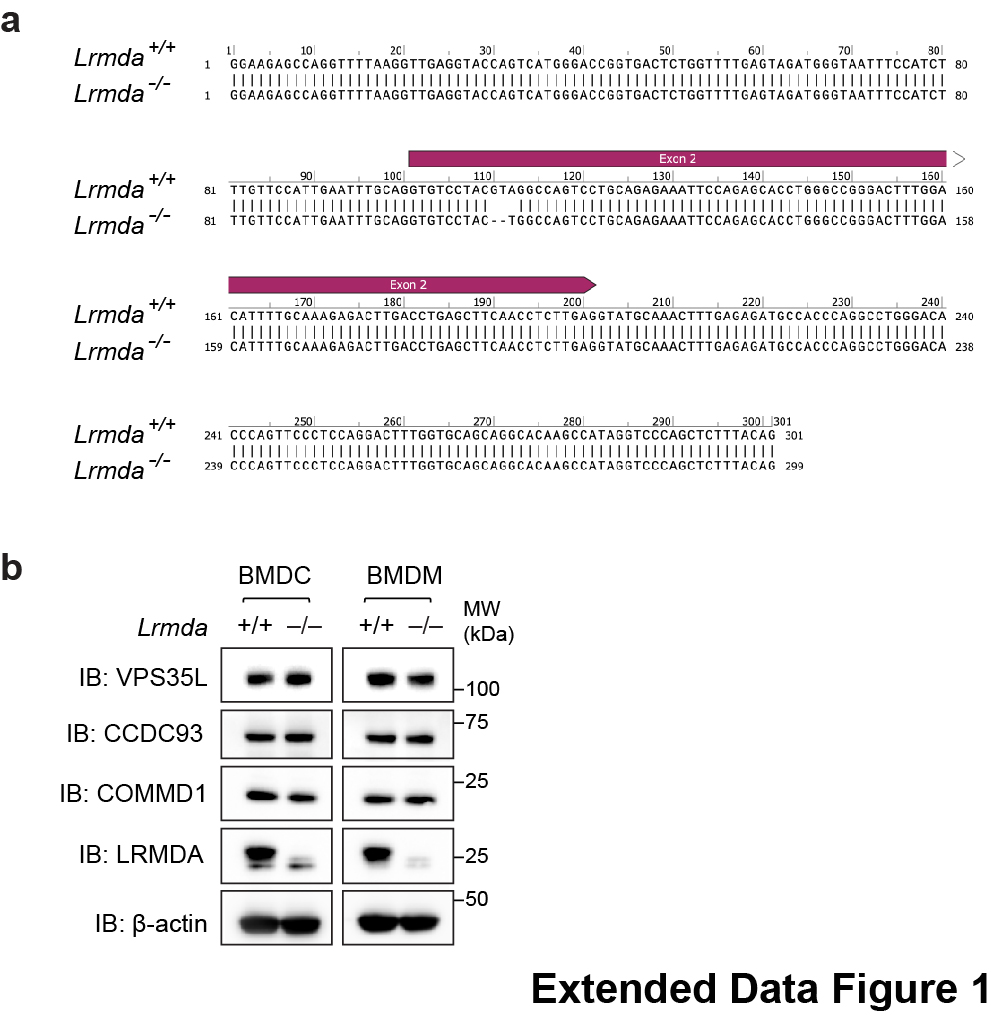

### Extended Data Figure 2

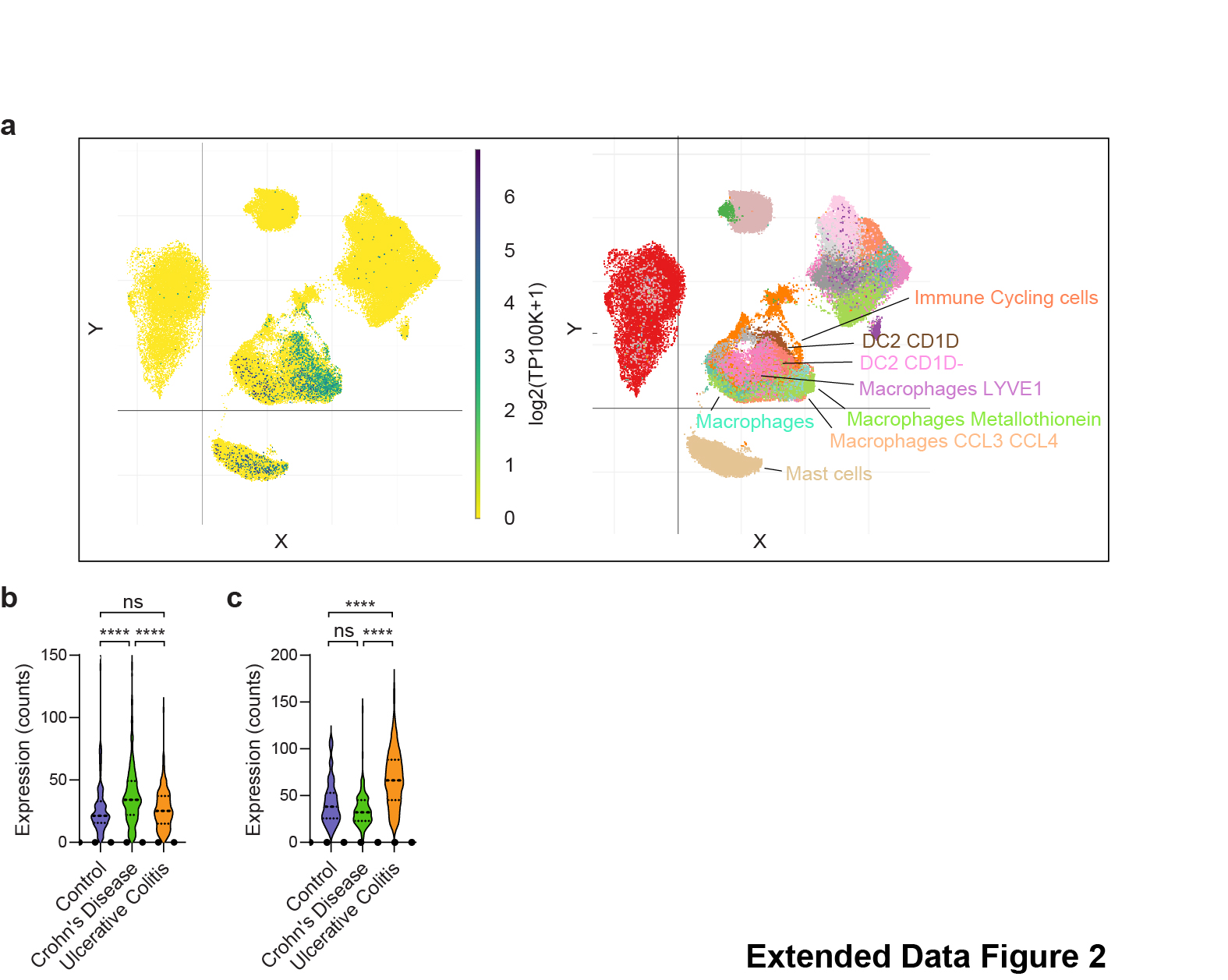

### Extended Data Figure 3

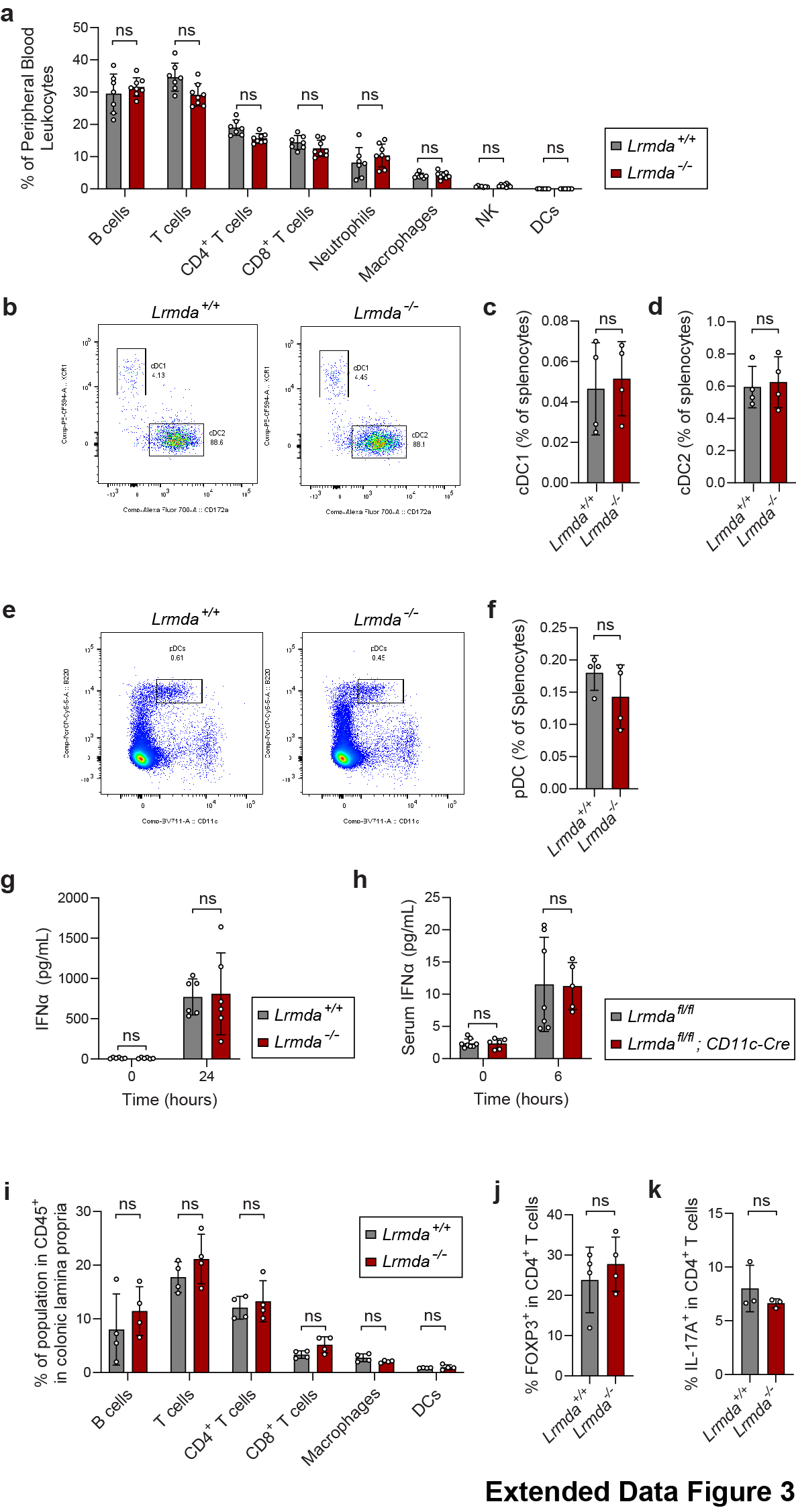
